# Activation of the actin/MRTF-A/SRF signalling pathway in pre-malignant mammary epithelial cells by P-cadherin is essential for transformation

**DOI:** 10.1101/2022.02.26.481995

**Authors:** Lídia Faria, Sara Canato, Tito T. Jesus, Margarida Gonçalves, Patrícia S. Guerreiro, Carla S. Lopes, Isabel Meireles, Eurico Morais de Sá, Joana Paredes, Florence Janody

## Abstract

Alterations in the expression or function of cell adhesion molecules have been implicated in all steps of tumour progression. Among those, P-cadherin expression is highly enriched in basal-like breast cancer, a molecular subset of triple-negative breast carcinomas, playing a central role in inducing cancer cell self-renewal, as well as collective cell migration and invasion capacity. To decipher the P-cadherin-dependent signalling network, we generated a humanised P-cadherin fly model, establishing a clinically relevant platform for functional exploration of P-cadherin effectors *in vivo*. We report that actin nucleators, MRTF and SRF are main effectors of P-cadherin functional effects. In addition, we validated these findings in a human mammary epithelial cell line with conditional activation of the Src oncogene, which recapitulates molecular events taking place during cellular transformation. We show that prior to triggering the gain of malignant phenotypes, Src induces a transient increase in P-cadherin expression levels, which correlates with MRTF-A accumulation, its nuclear translocation and the upregulation of SRF target genes. Moreover, knocking down P-cadherin, or preventing Factin polymerization with Latrunculin A, impairs SRF transcriptional activity. Furthermore, blocking MRTF-A nuclear translocation with CCG-203971 hampers proliferation, selfrenewal and invasion. Thus, in addition to sustaining malignant phenotypes, P-cadherin can also play a major role in the very early stages of breast carcinogenesis by promoting a transient boost of MRTF-A/SRF signalling through actin regulation.

## Introduction

Cell adhesion molecules (CAMs) are transmembrane receptor proteins widely expressed in epithelial tissues. Although their primary role is to maintain cell-to-cell contact and attachment to the extracellular matrix, they also integrate extracellular cues within cell intrinsic signalling, affecting the cytoskeleton organisation, intracellular responses and gene expression, and consequently cellular functions, such as cell growth, survival and invasion. It is therefore not surprising that alterations in their expression and/or activity influence malignant transformation, being considered potential targets for cancer therapy [1]. Yet, the mechanisms by which many of these CAMs impact cancer malignancy remain unclear.

Aberrant expression of the classical cell-cell adhesion molecule P-cadherin (P-cad) has been associated with aggressive tumour behaviour in breast, gastric, prostate, pancreatic, bladder and colorectal carcinomas among others [2]. In breast cancer, P-cad is overexpressed in a molecular subset of highly aggressive basal-like triple negative carcinomas, being significantly associated with a worse disease-free and overall patient survival [3]. P-cad promotes cell motility, collective cell migration and invasion capacity in breast cancer cells *in vitro* [4]. In addition, P-cad instructs cancer cells to acquire stem-like cell properties, thus contributing to the survival of aggressive breast cancer cells, and induces tumorigenic and metastatic capacity in *in vivo* breast cancer models [4–8]. Molecularly, P-cad activates α6β4 integrins and the nonreceptor tyrosine kinase Src, and interferes with E-cadherin (E-cad) function [6,8,9]. In addition, P-cad promotes GTPase-mediated signal transduction and affects the actin cytoskeleton [8,10,11]. Yet, how the actin cytoskeleton dynamics integrates into the P-cad-dependent signalling network remains to be established. Clarifying this network is fundamental to uncover potential therapeutic targets that could be used to counteract P-cad-mediated tumorigenic capacity.

Filamentous actin (F-actin) is assembled from monomeric actin subunits (G-actin). Polymerization occurs predominantly by extension of the fast-growing barbed ends of filaments, largely facilitated by the activity of diverse actin nucleators. Among those, the Arp2/3 complex, formed by seven subunits (Arp2, Arp3 and ARPC1-5), catalyses polymerization of new “daughter” filaments from the side of existing filaments to form branched networks. In contrast, formins generate the formation of linear, unbranched actin filaments, while Spire, a tandem-monomer-binding nucleator, recruits actin monomers to form polymerization seeds [12]. In addition to controlling key cellular processes that include the generation and maintenance of cell morphology and polarity, endocytosis, intracellular trafficking, contractility and cell division, the actin cytoskeleton is also a major regulator of signal transduction pathways [13]. One of those is the Myocardin-related transcription factor A (MRTF-A)/serum response factor (SRF) signalling pathway. MRTF-A contains a conserved N-terminus RPEL (arginine-proline-glutamine-leucine consensus sequence containing) domain that includes three actin-binding motifs (RPEL1, RPEL2, RPEL3), overlapping with an extended bipartite nuclear localization signal (NLS) [14,15]. In the absence of stimulation, MRTF-A localises in the cytoplasm due to its association with G-actin through the RPEL motifs [14]. This hinders the NLS of MRTF-A, preventing its recognition by the importin-α/β heterodimer, and thus, blocking its nuclear import [15]. Increased F-actin polymerization depletes the pool of G-actin which, consequently, dissociates from MRTF-A, allowing the access of import factors to the NLS to promote MRTF-A nuclear translocation, where it binds to SRF and activates the expression of target genes [15,16]. In turn, MRTF-A/SRF affect the actin cytoskeleton by regulating the expression of actin and of proteins controlling F-actin nucleation and organisation, explaining why the MRTF-A/SRF pathway plays a central role in the mobility of normal and cancer cells [17]. Indeed, increasing evidence indicates that MRTF-A has oncogenic properties, promoting proliferation, epithelial to mesenchymal transition (EMT), stemness abilities and metastasis [17–19].

Therapeutic approaches that would counteract the tumorigenic capacity of P-cad depend on a detailed understanding of its downstream signalling network. To this end, we generated a fly model with conditional expression of human P-cad, as the large arsenal of genetic tools in *Drosophila,* the rapid generation time and the high levels of conservation in genes encoding for cancer signalling pathways, allow powerful studies of underlying mechanisms [20]. We show that our humanised P-cad fly model is a clinically relevant platform for functional exploration of P-cad effectors *in vivo* and we identified *Drosophila* actin nucleators, MRTF and SRF, as suppressors of the P-cad-expressing wing phenotype. We have validated the role of the P-cad-dependent actin/MRTF-A/SRF signalling pathway in a human breast epithelial cell line, which recapitulates molecular events taking place during the transition from a normal to a malignant state. We provide evidence that P-cad plays a central role in the very early stages of breast carcinogenesis by promoting a transient boost of actin/MRTF-A/SRF signalling pathway activity, essential for the acquisition of pre-malignant and malignant phenotypes.

## Results

### Human P-cad is functional in *Drosophila* epithelia

*Drosophila* does not contain a *CDH3* ortholog [21]. Thus, to identify P-cad effectors relevant to its functional effect in cancer, we generated transgenic fly strains carrying human P-cad inducible with the Gal4-UAS system, which allows to control P-cad expression both temporally and spatially during fly development. To validate the humanised P-cad fly model, we searched for evidence that some P-cad-induced functional effects were recapitulated in fly epithelia. When mosaic expression was induced in the follicular epithelium, P-cad co-localized with the fly E-cad (DE-cad) apically, concentrating specifically at the cellular membranes juxtaposed to other P-cad expressing cells (Fig. 1A). Cross sections through the wing disc epithelium showed that P-cad expressed with the *nubbin-Gal4* (*nub-Gal4*) driver, also accumulated apically with DE-cad (Fig. 1B). Thus, human P-cad establishes *trans* junctional homophilic interactions in fly epithelia.

**Figure 1:**
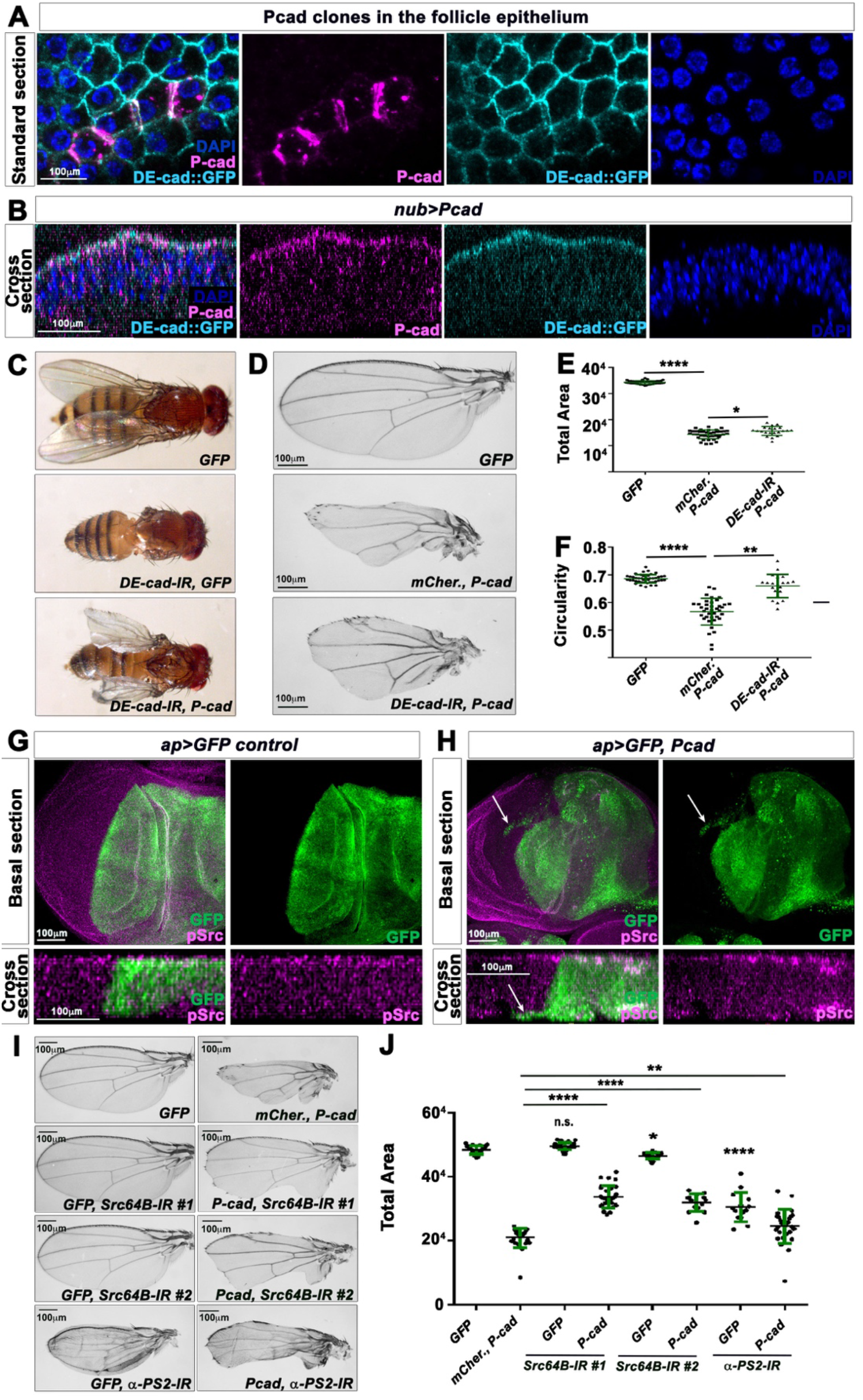
Human P-cad establishes cell-cell adhesion in *Drosophila* epithelia and affects wing development through DE-cad, Src64B and αPS2 integrin. **(A)** Standard confocal sections of P-cad-expressing clones in the follicle epithelium stained with DE-cad (cyan blue), P-cad (magenta) and DAPI (blue). **(B)** Cross section of third instar wing imaginal discs carrying a GFP knock-in into the DE-cad locus (cyan blue) and expressing *P-cad* under *nub*-Gal4 control, stained with P-cad (magenta) and DAPI (blue). **(C-D)** Adult **(C)** flies or **(D)** wings in which *nub*-Gal4 drives the indicated UAS-constructs. **(E-F)** Quantifications of the total wing area **(E)** and circularity **(F)** for the genotypes indicated. **(G-H)** Standard confocal sections (top panels) or cross section (bottom panels) of third instar wing imaginal discs expressing **(G)** *UAS-mCD8-GFP* (green) or **(H)** UAS-mCD8-*GFP* (green) and *UAS-P-cad* using the *ap*-Gal4 driver and stained with anti-phosphorylated Src (pSrc) (magenta). White arrows indicate cells migrating in the ventral compartment. **(I)** Adult wings in which *nub*-Gal4 drives the indicated UAS constructs. **(J)** Quantifications of the total wing area for the genotypes indicated. Error bars indicate SD; n.s. indicate non-significant; * indicates p<0.05; ** indicates p<0.01; **** indicates p<0.0001. Statistical significance was calculated using one-way ANOVA tests.

Since, P-cad has also been shown to compensate for the loss of E-cad in human cells [8,22], we asked whether human P-cad could substitute for DE-cad function. Indeed, while flies expressing a UAS-RNA interference (RNAi) construct against DE-cad in the wing epithelial primordium with *nub*-Gal4, failed to form wings, expressing P-cad in those animals partially restored wing development (Fig. 1C). Conversely, human E-cad is required for P-cad-induced tumour growth [8]. As P-cad-expressing wings were smaller, and failed to form a proper margin giving rise to a notched appearance, we asked whether reducing DE-cad function suppressed some aspects of this phenotype. Actually, knocking down DE-cad in these wings restored significantly their growth and circularity (Fig. 1D-F). Thus, in the fly wing epithelia, P-cad shows similar genetic interactions with DE-cad, as those reported with human E-cad in cancer cells [8,22].

In breast cancer cells, P-cad is also known to promote cell motility, collective cell migration and invasive capacities by potentiating the activation of the c-Src proto-oncogene and by signalling through c-Src and α6β4 integrins [4,6,9]. Thus, we tested if P-cad-expressing wing disc cells behave likewise. Whereas dorsal wing disc cells expressing GFP under *apterous*-Gal4 (*ap*-Gal4) maintained a straight boundary with GFP-negative ventral cells (Fig. 1G), those expressing P-cad with GFP suffered basal cell extrusion and migrated within the ventral compartment (see arrows in Fig. 1H). In addition, P-cad potentiated the levels of phosphorylated Src apically when compared to the ventral compartment, which has been used as control (Fig. 1H). We then tested if P-cad would signal through *Drosophila* Src and integrins. Expressing two independent UAS-RNAi against *Drosophila* Src64B had only marginal effects on wing size. However, both RNAi could drastically restore the growth defects of *nub>P-cad-expressing* wings (Fig. 1I, J). Moreover, knocking down the integrin αPS2 in P-cad-expressing wings also significantly restored wing growth, despite the penetrant inflated phenotype induced by reducing αPS2 function alone (Fig. 1I, J).

The reduced size of P-cad-expressing wing was not a consequence of tissue loss by apoptosis, as wing disc primordia expressing *P-cad* and *GFP* with *nub*-Gal4 only displayed low levels of activated Caspase 3 positive staining (Supp. Fig. 1A). Moreover, expressing the caspase inhibitor p35 did not restore the development of P-cad-expressing wing discs. Instead, these wings showed a worse phenotype, which could result from an additive effect between P-cad and p35, as p35 reduced wing size when expressed alone (Supp. Fig. 1B). Expressing P-cad in fast-dividing eye-antenna primordia, using the *eyeless*-Gal4 (*ey*-Gal4) driver, also drastically affected the development of the fly head (Supp. Fig. 1C). However, when expressed in postmitotic differentiating photoreceptor cells, using the GMR-Gal4, P-cad-expressing adult eyes did not display obvious defects, unlike adult eyes expressing *Rho1,* used as positive control (Supp. Fig. 1D). This suggests that the presence of P-cad specifically affects fast-dividing epithelia.

Taken together, we conclude that the consequences of expressing human P-cad in the fly wing epithelium are similar to those of overexpressing P-cad in breast cancer cells [6,8,9]. Thus the P-cad-dependent wing phenotype is suitable for the identification of P-cad effectors relevant to the acquisition of a tumorigenic phenotype.

### P-cad potentiates the activity of the DMRTF-DSRF signalling pathway

To identify P-cad effectors, we screened for signalling pathways suppressing the *nub>P-cad* adult wing phenotype when knocked down. Among the candidates recovered were components of the DMRTF-DSRF signalling pathway. Adult wings expressing RNAi against the *Drosophila* MRTF-A (*DMRTF-IR*) or the SRF blistered (*DSRF-IR*), together with UAS-*GFP* under *nub*-Gal4 control, did not display significant alterations in anterior-posterior length, but showed a slight reduction in total wing size area. However, both RNAi could significantly restore the defects in length and total area of *nub*>*P-cad* expressing wings (Fig. 2A). The DMRTF-DSRF signalling pathway is unlikely required to stabilise or localise P-cad, as P-cad levels were not significantly different in wing discs expressing *P-cad* and knocked down for *DMRTF* or *DSRF* compared to those expressing *P-cad* and *mCherry* (Fig. 2B). Moreover, cross sections through wing discs epithelia showed that P-cad localization was not altered by reducing DMRTF or DSRF function (Fig. 2C). To confirm that P-cad enhances the activity of the DMRTF/DSRF signalling pathway, we tested if P-cad was able to induce the nuclear translocation of a fusion between DMRTF and GFP (DMRTF.3XGFP), which was expressed using the ubiquitous *tubulin* promoter. We used the salivary gland of third instar larvae, as this monolayer tubular epithelium is composed of larger cells compared to those of the wing disc epithelium. When driven with the salivary gland specific driver *Sgs3-Gal4,* the levels of P-cad were very unequal between cells (Fig. 2D). DMRTF.3XGFP localised mainly in the cytoplasm in cells expressing low P-cad levels or in control cells carrying *Sgs3*-Gal4 only. In contrast, in cells expressing high P-cad levels, DMRTF.3XGFP levels were reduced in the cytoplasm and accumulated faintly in the nucleus. Quantification of the DMRTF.3XGFP intensity signal indicates that DMRTF.3XGFP accumulated significantly in the nucleus in P-cad-expressing cells (Fig. 2D). These observations are consistent with a role for P-cad in potentiating the DMRTF/DSRF signalling pathway. Accordingly, knocking down *MESK2 (MESK2-IR)* or *deterin* (*det*-IR), two DMRTF/DSRF target genes [23], restored the growth and circularity of P-cad-expressing wings (Supp. Fig. 2).

**Figure 2:**
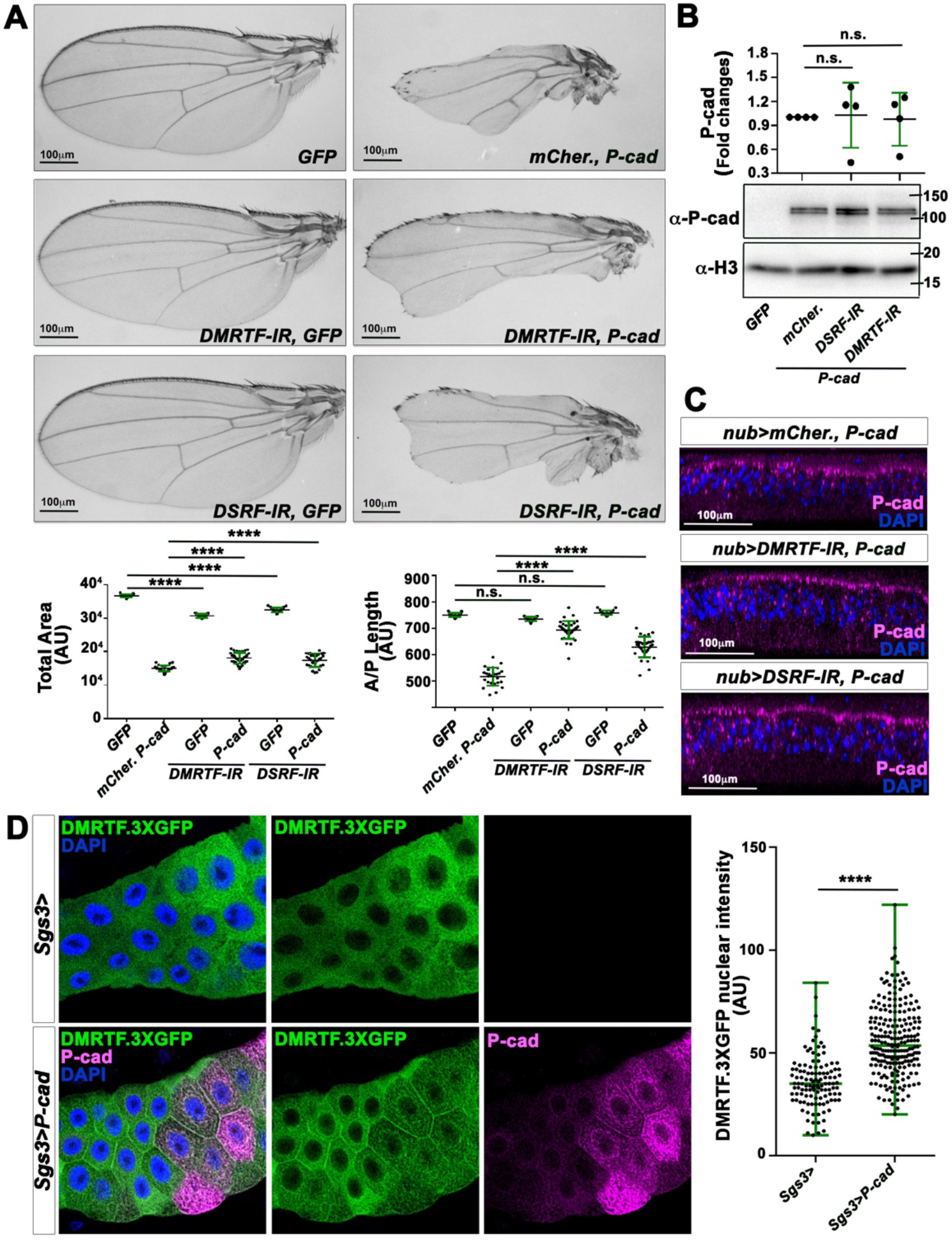
P-cad signals through the DMRTF-DSRF signalling pathway in the *Drosophila* wing. **(A)** (Top panels) adult wings in which*nub*-Gal4 drives the indicated UAS constructs. (Lower panels) quantifications of the total area (left) or anterior-posterior length (right) of adult wings for the genotypes indicated. **(B)** (Bottom panels) Western blots on protein extracts from wing imaginal discs expressing the indicated constructs under *nub*-Gal4 control, blotted with anti-P-cad and anti-Histone 3 (`α-H3`). (Top panel) quantification from four biological replicates of the ratio of P-cad levels normalised to α-H3 for the genotypes indicated. **(C)** Cross sections of third instar wing imaginal discs expressing the indicated constructs under *nub*-Gal4 control and stained with anti-P-cad (magenta) and DAPI (blue). **(D)** (Left panels) Standard confocal sections of third instar salivary glands, expressing *tub-DMRTF.3XGFP* (green) or *tub-DMRTF.3XGFP* and *UAS-P-cad* under *Sgs3-Gal4* control, stained with anti-P-cad (magenta) and DAPI (blue). (Right panel) quantification of the nuclear intensity signals of DMRTF.3XGFP for the genotypes indicated. Error bars indicate SD.; n.s. indicate non-significant; **** indicates p<0.0001. Statistical significance was calculated using one-way ANOVA tests.

Surprisingly, while P-cad enhanced DMRTF-DSRF signalling and reduced wing growth (Fig. 1E, 2A), the DMRTF-DSRF signalling pathway has been previously shown to enhance wing growth [24]. Indeed, adult wings homozygous for the *DSRF^2^* mutation were smaller. Conversely, overexpressing full length *DMRTF* enhanced wing growth (Supp. Fig. 3). On the other hand, expression of a constitutively active form of *DMRTF* (*DMRTF-ΔN*), with deletion of the N terminus, which renders nuclear and active mammalian and *Drosophila* MRTF [25,26], reduced wing size and circularity (Supp. Fig. 3), reminiscent of the P-cad functional’s effect. These observations suggest that low increase in DMRTF/DSRF activity promotes tissue growth, while higher activity affects wing growth and differentiation. Taken together, our observations are consistent with a role of P-cad in promoting high DMRTF/DSRF signalling activity.

### P-cad signals through the actin cytoskeleton

As G-actin levels regulate the nucleus–cytoplasm shuttling of MRTF-A [15,16], we asked if P-cad expression would affect the actin cytoskeleton. Accordingly, expressing *P-cad* and *GFP* with *ap*-Gal4 in the dorsal wing disc epithelium, increased the pool of F-actin, which accumulated mainly on the basal surface of the disc epithelium (Fig. 3, compare B with A). We then tested if reducing F-actin nucleation could suppress the P-cad-expressing wing phenotype. Indeed, knocking down components of the Arp2/3 complex or the tandem monomer binder *spire*, or even the formin *diaphanous* (*dia*), significantly restored the anterior-posterior length and size of *nub>P-cad*-expressing wings (Fig. 3C and Supp. Fig. 4). To test the possibility that the actin cytoskeleton rescues the P-cad wing phenotype by acting upstream of P-cad, we analysed the effect of knocking down these actin nucleators on P-cad levels. Knocking down *arpc2* or *spire* significantly reduced P-cad levels (Fig. 3D). In contrast, the levels of P-cad were not significantly different in wing disc extracts expressing *P-cad* and *dia-IR,* compared to those expressing *P-cad* and *mCherry* (Fig. 3E). Moreover, P-cad remained apically localised in wing disc epithelia expressing *dia-IR*, further suggesting that Dia is required downstream of P-cad to affect wing development. Although Arpc2 and Spire regulate P-cad protein stability, they could have additional functional effects downstream of P-cad. Consistent with this hypothesis, knocking down DMRTF or DSRF did not further suppress the defects of wings expressing *P-cad* and *arpc2-IR* or *spire-IR* (Supp. Fig. 5A). As the MRTF-SRF pathway is well known to control the expression of many genes related to the actin cytoskeleton [27], we also tested if Arpc2 or Spire promotes P-cad functional effects downstream of the MRTF/SRF pathway. However, knocking down *arpc2* or *spire* did not rescue the wing phenotype induced by expressing *MRTFΔN*. On the contrary, expressing *arpc2-IR* or *spire-IR* further reduced the size or anterior-posterior length of these wings (Supp. Fig. 5B). Taken all together, these observations suggest that the actin cytoskeleton, controlled by the Arp2/3 complex and Spire, stabilises P-cad. In addition, actin regulation by the Arp2/3 complex, Spire and Dia is involved downstream of P-cad, likely by enhancing the activity of the DMRTF/DSRF signalling pathway.

**Figure 3:**
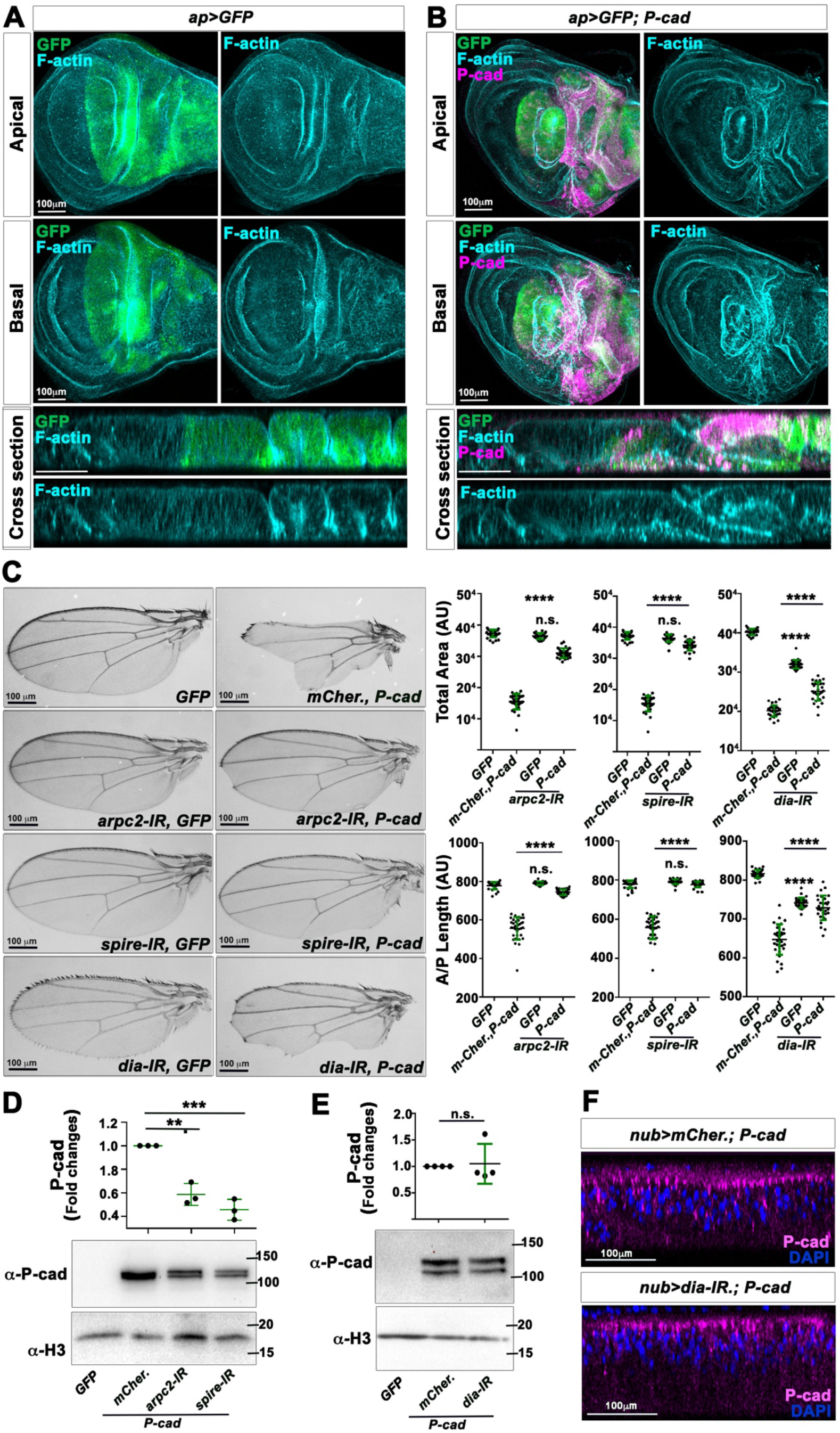
Actin nucleation is required for P-cad functional effect in the fly wing. **(A,B)** Standard confocal sections of the apical (top panel) or basal (middle panel) surfaces of third instar wing imaginal discs or cross sections through wing imaginal discs (bottom panel), expressing **(A)** *UAS-mCD8-GFP* (green) or **(B)** *UAS-mCD8-GFP* (green) and *UAS-P-cad* under *ap*-Gal4 control and stained with Phalloidin (cyan blue) to mark the actin cytoskeleton and anti-P-cad (magenta). **(C)** (Left panels) Adult wings in which *nub*-Gal4 drives the indicated UAS-constructs. (Right panels) Quantifications of the total area (top) or anterior-posterior (A/P) length (bottom) of adult wings for the genotypes indicated. **(D)** (Bottom panels) Western blots on protein extracts from wing imaginal discs expressing the indicated UAS-constructs under *nub*-Gal4 control, blotted with anti-P-cad and anti-Histone 3 (`α-H3`). (Top panel) Quantification from three biological replicates of the ratio of P-cad levels normalised to α-H3 for the genotypes indicated. **(E)** (Bottom panels) Western blots on protein extracts from wing imaginal discs expressing the indicated UAS-constructs under *nub*-Gal4 control, blotted with anti-P-cad and anti-Histone 3 (`α-H3`). (Top panel) Quantification from four biological replicates of the ratio of P-cad levels normalised to α-H3 for the genotypes indicated. **(F)** Cross sections of third instar wing imaginal discs expressing the indicated UAS-constructs under *nub*-Gal4 control and stained with anti-P-cad (magenta) and DAPI (blue). Error bars indicate SD; n.s. indicate non-significant; ** indicates p<0.01; *** indicates p<0.001; **** indicates p<0.0001. Statistical significance was calculated using one-way ANOVA tests.

### P-cad transiently accumulates in TAM-treated MCF10A-ER-Src cells

To validate the role of the MRTF-A/SRF signalling pathway in promoting tumorigenic phenotypes downstream of P-cad in a human context, we used the mammary epithelial cell line MCF10A-ER-Src, which contains a fusion between the viral c-Src kinase orthologue and the ligand-binding domain of the oestrogen receptor (ER), inducible with tamoxifen (TAM) treatment [28,29]. cDNA microarrays analysis indicates that TAM-treated cells upregulate *CDH3* (P-cad codifying gene) during the transition from normal to transformed cells, which involves the gain of sustained proliferative abilities 12 hours after TAM treatment, EMT and the acquisition of stem-like properties 24 hours after treatment, and invading abilities 36 to 45 hours after TAM treatment [30–33]. We have reported that the gain of self-sufficiency in growth properties and the progression towards a fully transformed state, result from a transient increase in actomyosin-dependent cell stiffening during the first 12 hours of TAM treatment, which is associated with a transitory accumulation of F-actin (Fig. 4A) [31]. We therefore checked if this early accumulation of F-actin was associated with P-cad upregulation. The ratio of P-cad mRNA levels between cells treated with TAM and EtOH indicated that P-cad levels were significantly upregulated by 2.13 fold, 2 hours after TAM treatment, maintained significant higher expression up to 12 hours after treatment and progressively dropped to initial levels 24 and 36 hours after treatment (Fig. 4B). This transient increase in P-cad mRNA levels was translated into an increase at protein levels in TAM-treated cells, compared to those treated with EtOH for the same time (Fig. 4C). Moreover, quantification of the Median Fluorescence Intensity (MFI) of membrane-associated P-cad by Fluorescent Activating Cell sorting (FACS) showed that the levels of P-cad were significantly increased between 6 and 24 hours after TAM treatment (Fig. 4D). Strikingly, a sub-population of MCF10A-ER-Src cells were highly enriched in membrane-associated P-cad, starting 6 hours after TAM treatment, compared to those treated with EtOH (Fig. 4E). Thus, P-cad accumulation correlates with the transient accumulation of F-actin in pre-malignant MCF10A-ER-Src cells.

**Figure 4:**
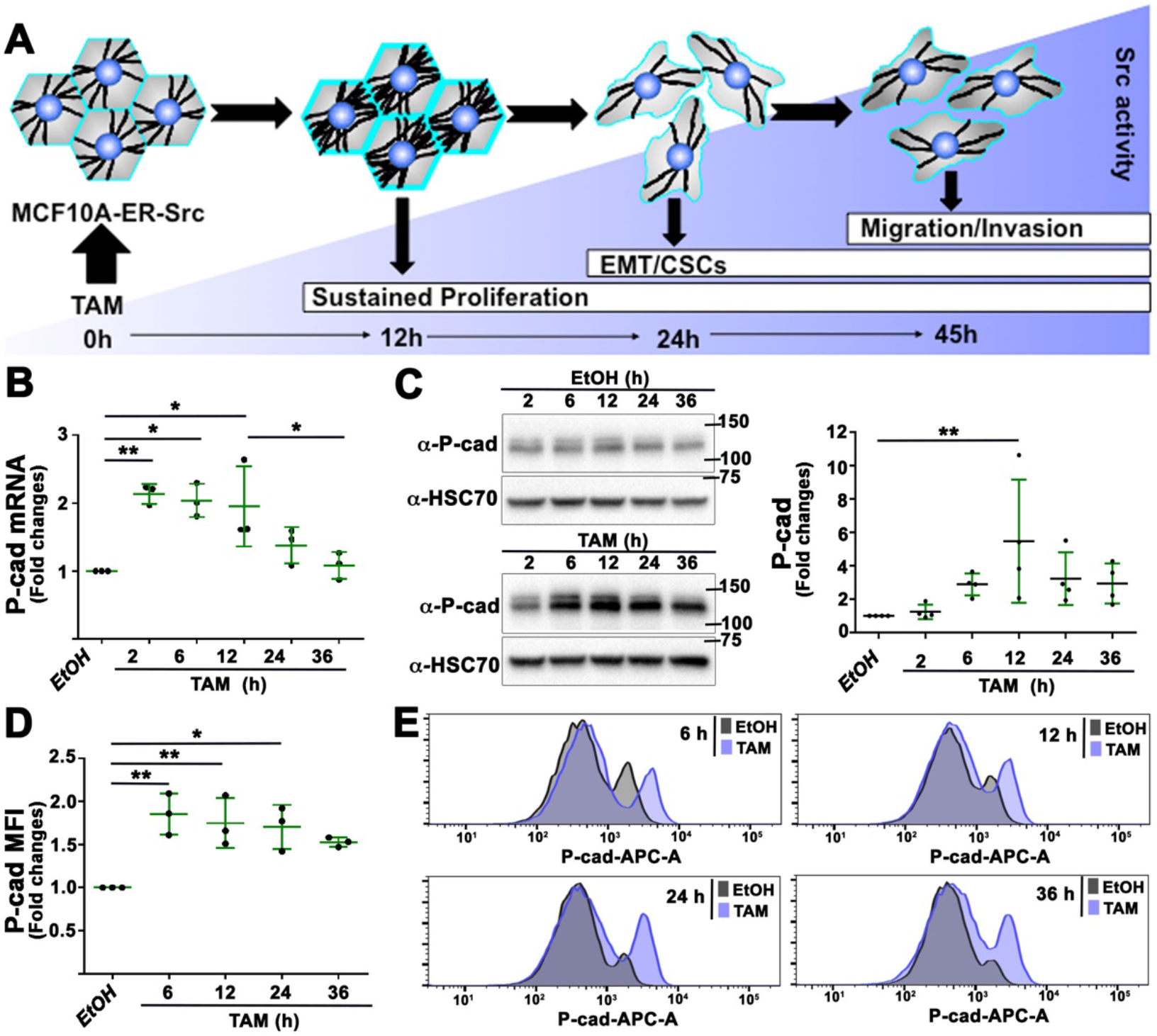
P-cad transiently accumulates in TAM-treated MCF10A-ER-Src cells. **(A)** Schematic of the series of events taking place during transformation of the MCF10A-ER-Src cell line treated with TAM. MCF10A-ER-Src cells transiently accumulate F-actin-stress fibres 12 hours after TAM treatment. In turn, stress fibre organisation sustains cell proliferation, as well as further potentiates Src activity. Cells then undergo EMT and acquire CSC properties 24 hours after TAM treatment and gain invading abilities at 45 hours of treatment [30–32]. **(B)** *P-cad* mRNA levels in extracts from MCF10A-ER-Src cells treated with TAM for 2, 6, 12, 24 and 36 hours, normalised to those from MCF10A-ER-Src cells treated with EtOH for the same time points. **(C)** (Left panels) Western blots on protein extracts from MCF10A-ER-Src cells treated with EtOH (upper panels) or TAM (lower panels) for 2, 6, 12, 24 or 36 hours, blotted with anti-P-cad or anti-HSC70. (Right panel) Ratio of P-cad levels between TAM- and EtOH-treated MCF10A-ER-Src cells for the same time points, normalised to HSC70. **(D)** Median Fluorescence Intensity (MFI) of membrane-associated P-cad in MCF10A-ER-Src cells treated with TAM for 6, 12, 24 and 36 hours, normalised to those from MCF10A-ER-Src cells treated with EtOH for the same time points. **(E)** Histograms of membrane-associated P-cad in MCF10A-ER-Src cells treated with TAM or EtOH for 6, 12, 24 and 36 hours. Error bars indicate SD; * indicates p<0.05; ** indicates p<0.01. Statistical significance was calculated using unpaired t-test when comparing 2 conditions or one-way ANOVA tests in case of multiple comparisons.

### P-cad triggers MRTF-A/SRF signalling in TAM-treated MCF10A-ER-Src cells

To determine if the transient accumulation of F-actin and P-cad was also associated with a boost of SRF transcriptional activity, we measured the activity of a SRF luciferase reporter, which contains a luciferase construct controlled by three SRF binding sites [34]. Temporal analysis showed that Luciferase activity rose up to 3.8 fold during the first 6 hours of TAM treatment, before dropping 36 hours after treatment to levels similar to those of EtOH treated cells (Fig. 5A). This increase in SRF transcriptional activity was also associated with the transient accumulation of MRTF-A by Western Blot during the first 24 hours of TAM treatment (Fig. 5B), as well as with the nuclear accumulation of MRTF-A in MCF10A-ER-Src cells treated with TAM for 6 hours (Fig. 5C). Furthermore, the expression of SRF, which has been previously identified as a direct MRTF-A/SRF target gene [35], was also transiently upregulated 2 to 6 hours after TAM treatment (Fig. 5D).

**Figure 5:**
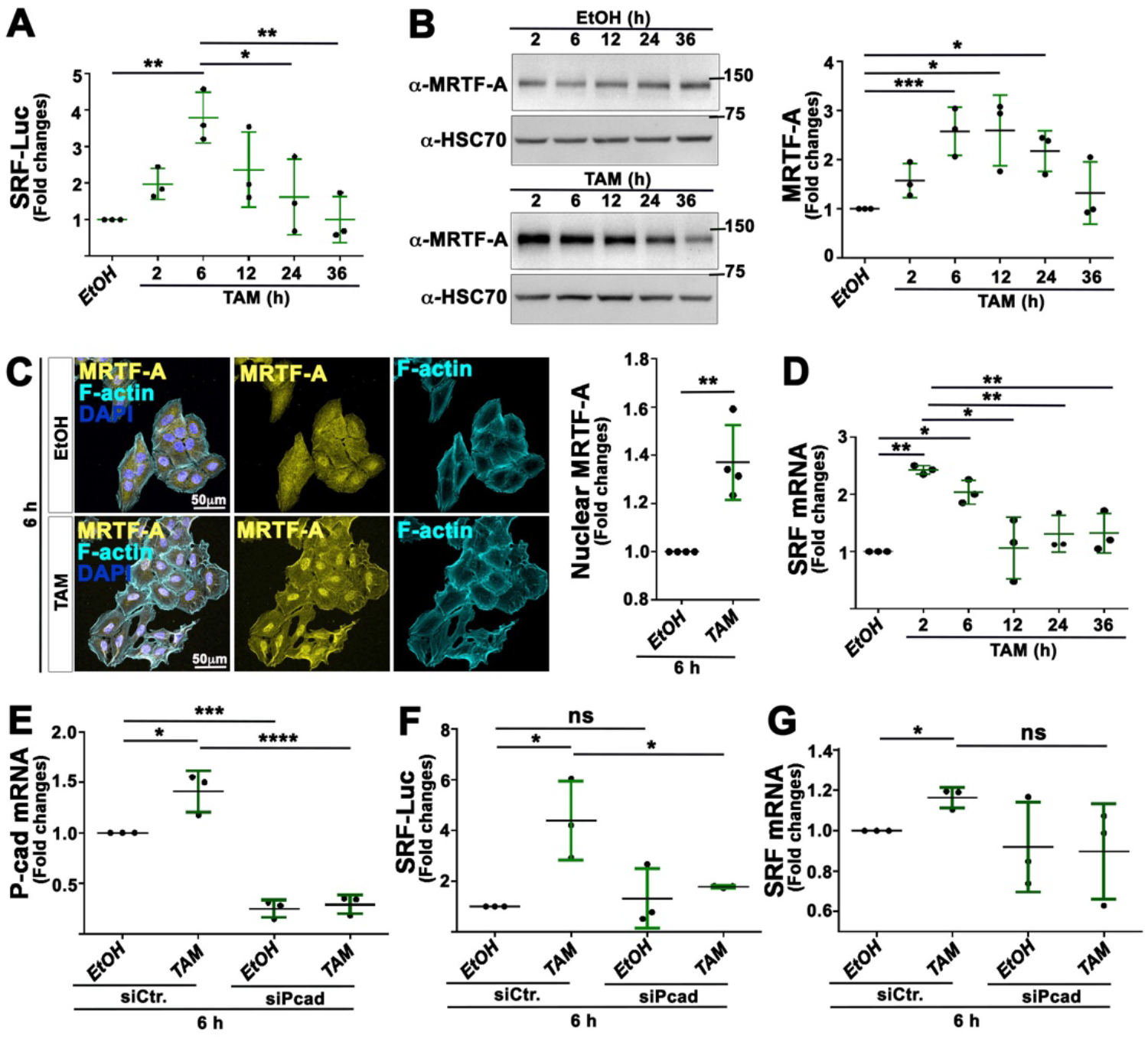
Knocking down P-cad prevents the transient increase in SRF transcriptional activity in TAM-treated MCF10A-ER-Src cells. **(A)** Fold changes in SRF Luciferase activity between MCF10A-ER-Src cells treated with EtOH or TAM, transfected with the SRF-responsive-Luc reporter gene. **(B)** (Left panels) Western blots on protein extracts from MCF10A-ER-Src cells treated with EtOH or TAM for 2, 6, 12, 24 or 36 hours, blotted with anti-MRTF-A or anti-HSC70. (Right panel) Ratio of MRTF-A levels between TAM- and EtOH-treated MCF10A-ER-Src cells for the same time points, normalised to HSC70. **(C)** (Left panels) Standard confocal sections of MCF10A-ER-Src cells treated with EtOH or TAM for 6 hours, stained with Phalloidin (cyan blue), anti-MRTF-A (yellow) and DAPI (blue). (Right panel) Quantification of the intensity of MRTF-A in the nucleus in MCF10A-ER-Src cells treated with EtOH or TAM for 6 hours. **(D)** *SRF* mRNA levels in extracts from MCF10A-ER-Src cells treated with TAM for 2, 6, 12, 24 and 36 hours, normalised to those from MCF10A-ER-Src cells treated with EtOH for the same time points. **(E)** *CDH3* mRNA levels on extracts from MCF10A-ER-Src cells expressing *siCtr*. or *siP-cad* and treated with EtOH or TAM for 6 hours. **(F)** Fold changes in SRF Luciferase activity between MCF10A-ER-Src cells transfected with the SRF-responsive-Luc reporter gene, expressing *siCtr*. or *siP-cad* and treated with EtOH and TAM for 6 hours. **(G)** *SRF* mRNA levels on extracts from MCF10A-ER-Src cells expressing *siCtr*. or *siP-cad* and treated with EtOH or TAM for 6 hours. Error bars indicate SD; * indicates p<0.05; ** indicates p<0.01; *** indicates P<0.001. Statistical significance was calculated using unpaired t-test when comparing 2 conditions or one-way ANOVA tests in case of multiple comparisons.

We then knocked down P-cad using small interfering RNA (*siPcad*) to test if the transient increase in SRF transcriptional activity after TAM treatment required P-cad. MCF10A-ER-Src cells transfected with *siPcad* and treated with EtOH or TAM for 6 hours showed a 75% and 79% reduction, respectively of *CDH3* mRNA levels, compared to EtOH or TAM-treated cells transfected with a siRNA control *(siCtr)* (Fig. 5E). Knocking down P-cad in TAM-treated cells significantly abrogated the transient increase in SRF-dependent Luciferase activity (Fig. 5F). SRF mRNA levels were also reduced upon P-cad silencing, although this decrease was not statistically significant (Fig. 5G). Taken all together, we conclude that the transient accumulation of P-cad in TAM-treated MCF10A-ER-Src cells, triggers a temporal increase in MRTF-A/SRF signalling prior to the acquisition of malignant features.

### Actin polymerization induces SRF activity in TAM-treated MCF10A-ER-Src cells

TAM treatment promotes a transient increase in actin polymerization in MCF10A-ER-Src cells within the first 12 hours [31]. Thus, we probed if the increased MRTF-A/SRF signalling activity depended on transient F-actin accumulation, by treating cells with Latrunculin A (LatA), which blocks the dissociation of the G-actin:MRTF-A complex [25,36]. LatA drastically reduced F-actin levels in MCF10A-ER-Src cells treated with either EtOH or TAM for 12 hours (Fig. 6A), but did not affect cell viability, which was maintained high to about 96 to 97% (Fig. 6B). While LatA treatment had no significant effect on SRF Luciferase activity in MCF10A-ER-Src cells treated with EtOH for 6 hours, it abrogated the Luciferase activity of cells treated with TAM for 6 hours (Fig. 6C). Thus, actin polymerization is required to induce the transient increase in SRF signalling activity.

**Figure 6:**
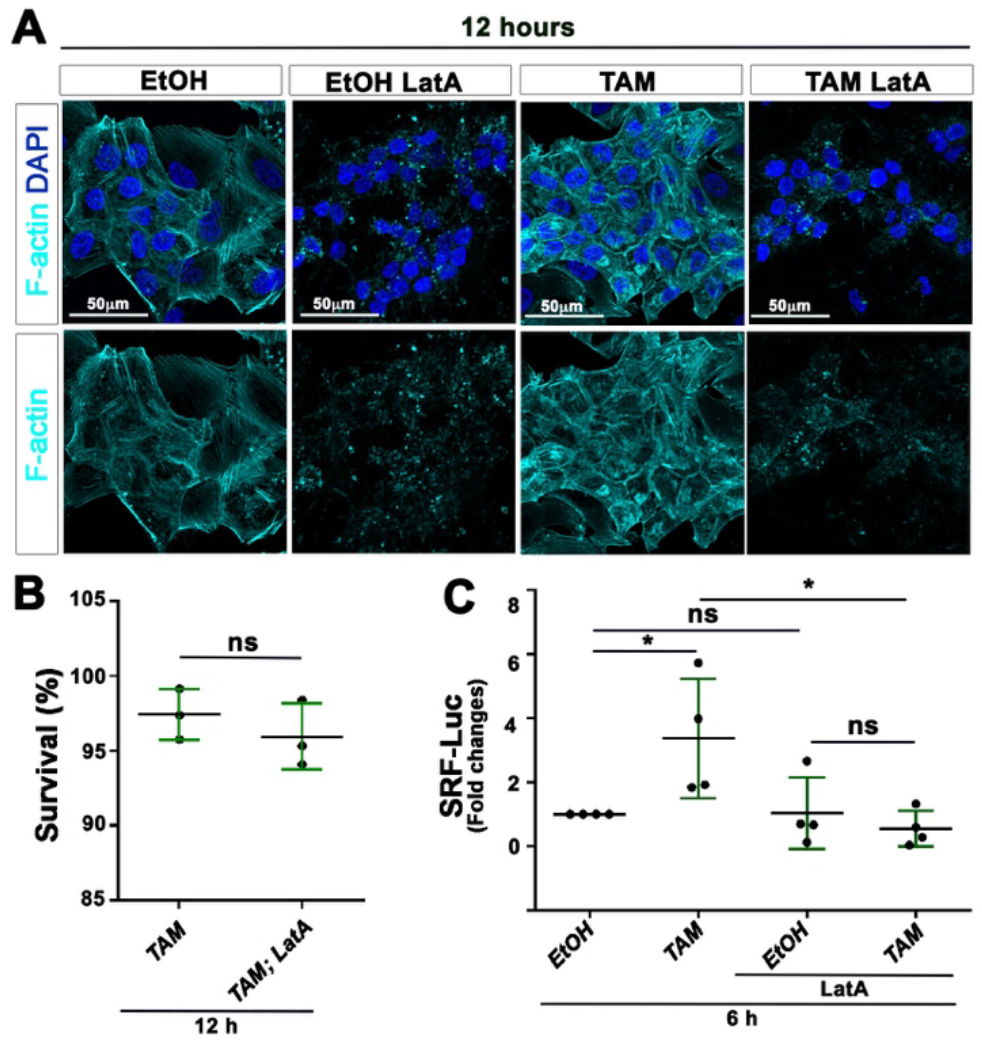
LatA inhibits SRF transcriptional activity in MCF10A-ER-Src cells treated with TAM for 6 hours. **(A)** Standard confocal sections of MCF10A-ER-Src cells treated with EtOH or TAM for 12 hours, in the presence or absence of LatA, stained with Phalloidin (cyan blue) to mark F-actin and DAPI (blue). Scale bars represent 50 μm. **(B)** Percentage of viable MCF10A-ER-Src cells treated with TAM for 12 hours, in the presence or absence of LatA. **(C)** Fold changes in Luciferase activity between MCF10A-ER-Src cells treated with EtOH or TAM for 6 hours in the presence or absence of LatA, transfected with the SRF-responsive-Luc reporter gene. Error bars indicate SD; n.s. indicate non-significant; * indicates P<0.05; Statistical significance was calculated using unpaired t-test when comparing 2 conditions or one-way ANOVA tests in case of multiple comparisons.

### MRTF-A triggers malignant transformation in TAM-treated MCF10A-ER-Src cells

To explore the role of the transient increase in MRTF-A/SRF signalling activity prior to the acquisition of malignant features, we used the small molecule inhibitor CCG-203971, which has been reported to block the nuclear localization and activity of MRTF-A, without causing cellular toxicity [37,38]. We first confirmed that the expression of the minimal SRF-dependent Luciferase reporter gene was dependent on MRTF-A activity. Accordingly, MCF10A-ER-Src cells co-treated with CCG-203971 and TAM failed to increase SRF-dependent Luciferase activity, compared to those that were treated with TAM only (Fig. 7A). We then tested the consequences of blocking MRTF-A activity on the ability of TAM-treated MCF10A-ER-Src cells to acquire transformed features. In the absence of Epidermal Growth Factor (EGF) and low serum concentration, TAM-treated MCF10A-ER-Src cells showed a significant increase in the percentage of cells in S-phase of the cell cycle compared to EtOH-treated cells, 12 and 24 hours after treatment. MRTF-A activity is required for TAM-treated cells to gain self-sufficiency in growth properties, as CCG-203971 abrogated their proliferative advantage 12 and 24 hours after TAM treatment (Fig. 7B). As previously reported, MCF10A-ER-Src cells treated with TAM for 36 hours formed significantly more mammospheres, compared to those treated with EtOH. Blocking MRTF-A activity using CCG-203971 drastically reduced the mammosphere-forming ability of TAM-treated cells (Fig. 7C), indicating that MRTF-A activity is required for TAM-treated cells to acquire self-renewing abilities. We then accessed the effect of CCG-203971 on the ability of TAM-treated MCF10A-ER-Src cells to invade in collagen. While MCF10A-ER-Src spheroids treated with EtOH for 36 hours, maintained a round shape with well-defined borders, those that were treated with TAM for the same time displayed irregular borders with cells invading into the collagen matrix (Fig. 7D). CCG-203971 reduced the number of invading TAM-treated spheroids and restored their circularity. Altogether, these observations suggest that the P-cad-dependent increase of MRTF-A activity provides proliferative abilities to TAM-treated MCF10A-ER-Src cells and is required for malignant transformation.

**Figure 7:**
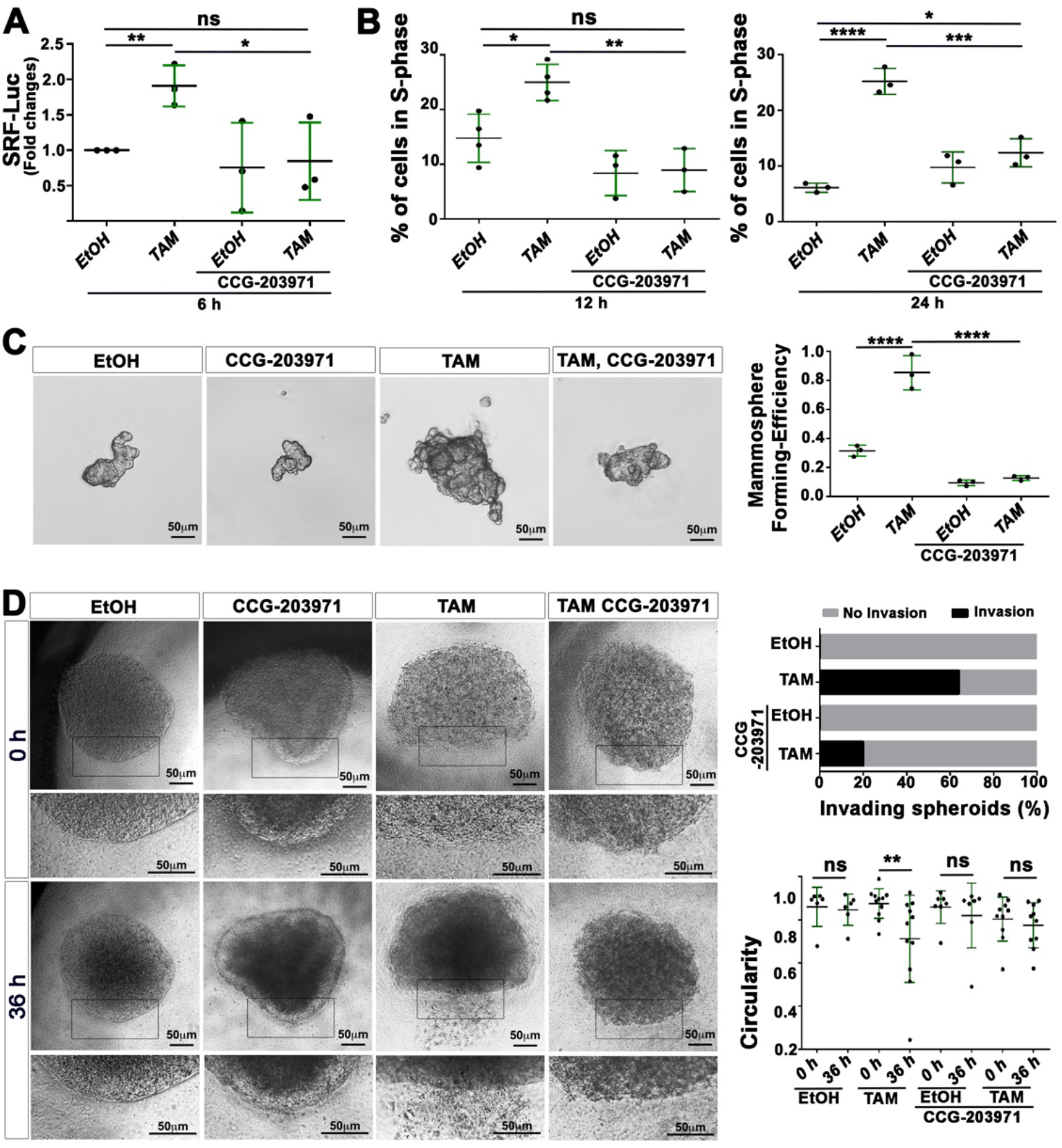
CCG-203971 reduces the ability of TAM-treated MCF10A-ER-Src cells to sustain proliferation, to grow mammospheres and to invade in collagen. **(A)** Fold changes in Luciferase activity between MCF10A-ER-Src cells treated with EtOH or TAM, in the presence or absence of CCG-203971, transfected with the SRF-responsive-Luc reporter gene. **(B)** Percentage (%) of MCF10A-ER-Src cells in S-phase, treated for 12 hours (Left panel) or 24 hours (Right panel) with EtOH or TAM in the presence or absence of CCG-203971. **(C)** (Left panel) Representative images of MCF10A-ER-Src mammospheres treated with EtOH or TAM, in the presence or absence of CCG-203971. Scale bars represent 50 μm. (Right panel) Mammosphere-forming efficiency of MCF10A-ER-Src mammospheres treated with EtOH or TAM, in the presence or absence of CCG-203971. **(D)** (Left panel) Representative images of MCF10A-ER-Src spheroids in collagen before treatment (0 h) or after 36 hours (36 h) of treatment with EtOH or TAM, in the presence or absence of CCG-203971. The lower panels are magnifications of the region delimited by the black line in the middle panel. Scale bars represent 50 μm. (Upper right panel) Percentage (%) of invading MCF10A-ER-Src spheroids treated with EtOH or TAM, in the presence or absence of CCG-203971. (Lower right panel) Circularity of MCF10A-ER-Src spheroids in collagen before treatment (0 h) or after 36 hours (36 h) of treatment with EtOH or TAM, in the presence or absence of CCG-203971. Error bars indicate SD.; * indicates P<0.05; ** indicates p<0.01; *** indicates p<0.001; **** indicates P<0.0001. Statistical significance was calculated using unpaired t-test in panel D and one-way ANOVA tests in panel A, B and C.

## Discussion

Here, we provide evidence that the humanised P-cad fly model we have generated is a clinically relevant platform for the identification of P-cad effectors. In agreement with observations reported in human cells [39], we show that P-cad also establishes *trans* junctional homophilic interactions when expressed in *Drosophila* epithelia. Moreover, P-cad induces cell invasion and signals through DE-cad, Src and Integrin in the wing disc epithelium, similar to its functional effect in breast cancer cells [6,8,9]. Furthermore, P-cad expression affects specifically undifferentiated/progenitor cells of the eye and wing discs, without impacting on post-mitotic epithelial cells fully engaged in the differentiation pathway. Consistent with these observations, P-cad promotes the maintenance of an undifferentiated state in the normal mammary gland, but has no apparent effect when expressed in differentiated mammary epithelial cells [40,41]. Yet, our P-cad fly model does not mimic all outcomes caused by P-cad in breast cancer cells. In these cells, P-cad promotes tumorigenic capacity [8]. In contrast, when expressed in the proliferating wing or eye disc primordium, P-cad prevents tissue growth. These distinct P-cad-dependent functional outcomes can result from cellular and tissue-specific responses. Accordingly, in hepatocarcinoma cells, P-cad has been shown to restrict cell proliferation [42].

Our humanised fly model uncovered DMRTF and DSRF, as modifiers of the wing phenotype produced by P-cad overexpression. Like P-cad, expressing a constitutively active form of *DMRTF* reduces wing growth. As P-cad levels or localization are not affected by knocking-down DMRTF and DSRF, and since P-cad induces DMRTF nuclear translocation, we propose that the MRTF-A/SRF axis is a key P-cad effector pathway in human cells. We have validated this hypothesis using a mammary epithelial cell line, which recapitulates molecular events taking place during the transition from a normal to a malignant state. While P-cad overexpression has been implicated in sustaining malignant phenotypes in fully transformed breast cancer cells [2], our observations suggest that P-cad could also play a major role in the very early stages of breast carcinogenesis by promoting a transient boost of MRTF-A/SRF signalling pathway activity, essential for the acquisition of pre-malignant and malignant phenotypes. We show that one of the first responses to Src activation in MCF10A-ER-Src cells is the transient accumulation of P-cad, which correlates with MRTF-A accumulation, its nuclear translocation and the upregulation of SRF itself and of a minimum SRF transcriptional reporter gene. Moreover, knocking down P-cad or preventing MRTF-A nuclear translocation with CCG-203971 [37,38], impedes SRF transcriptional activity. Furthermore, we show that the transient boost of MRTF-A activity provides proliferative advantages, stemness and invading abilities to TAM-treated MCF10A-ER-Src cells. Consistent with our observations, overexpressing MRTF-A in untransformed MCF10A acini is sufficient to induce pre-malignant spheroid formation caused by the dysregulation of cell cycle genes and EMT [18]. This mechanism may also be in place in other cancer subtypes, including those of the pancreas, as P-cad has tumour-promoting effects in these cancer cells, and the overexpression of MRTF-A in normal pancreatic cells induces stem cells features and EMT [43,44]. In addition to signal through SRF, MRTF-A could also promote cellular transformation independently of SRF, through direct binding to DNA [45]. Consistent with this possibility, microarray analysis indicates that tenascin C, whose expression is regulated directly by MRTF-A, is upregulated in TAM-treated MCF10A-ER-Src cells [33,45]. Moreover, MRTF-A triggers the expression of a specific set of pro-proliferative genes, independently of SRF [46]. MRTF-A activity could, in turn, regulate P-cad expression, whose promoter contains single nucleotide polymorphism (SNP) in the SRF CArG binding site [47]. Although activation of the P-cad/actin/MRTF-A/SRF pathway is a transient event during the early stages of cellular transformation, a small population of TAM-treated MCF10A-ER-Src cells could maintain high P-cad-dependent MRTF-A/SRF activity, as some cells retain high levels of membrane-associated P-cad 24 and 36 hours after TAM treatment. Consistent with a role of MRTF-A/SRF signalling in mediating P-cad functional effects in fully transformed cells, MRTF-A/SRF has been shown to play an important role in the metastatic lung colonisation of breast cancer cells and in drug resistance of basal cell carcinoma [48,49].

We have previously reported that MCF10A-ER-Src cells undergo a transient increase in Factin assembly, as well as show actomyosin-dependent cell stiffening during the first 12 hours of TAM treatment. We have shown that preventing cell stiffening, without interfering with Factin accumulation, prevents ERK-dependent cell proliferation, reduces Src activation and restricts the progression towards a fully transformed state [31]. We now provide evidence that the transient increase in F-actin assembly, triggered in part by P-cad, induces MRTF-A/SRF-dependent malignant transformation. First, P-cad causes F-actin accumulation in the wing disc epithelium. Accordingly, P-cad affects the actin cytoskeleton in breast cancer cells [9,11]. Second, P-cad requires the actin-nucleating activity of the Arp2/3 complex, Spire and Dia to affect wing development. Third, blocking the dissociation of the G-actin:MRTF-A complex using LatA [25,36], impedes SRF transcriptional activity in TAM-treated cells. The increase in actomyosin-dependent cell stiffening could also enhance MRTF-A activity, as MRTF-A has been shown to respond to mechanical stress [18,50,51]. P-cad could promote actin nucleation through activation of Rho-GTPases, like Rac1 and Cdc42, which are well-known regulators of MRTF-A nuclear translocation and SRF transcription [17,25]. Thus, during the early stage of breast cancer, major alterations of the actin cytoskeleton could control signalling pathways through distinct mechanisms: an increase in F-actin assembly would induce signalling pathways, such as MRTF-A/SRF, while actomyosin activity would trigger the mechanical induction of signalling components, such as ERK and Src, highlighting the key contribution of actin cytoskeleton regulation to carcinogenesis.

## Acknowledgements

Stocks obtained from the Bloomington Drosophila Stock Center (NIH P40OD018537) were used in this study. We specially thank K. Struhl and G. Posern for reagents, the IGC fly, imaging, cytometry and genomics facilities for technical assistance, and S. Tavares for comments on the manuscript. We thank all members of F.J.’s and J.P. labs for helpful discussions. We acknowledge the Cell Culture and Genotyping, the Genomics, the Translational Cytometry, the Bioimaging and the Advanced Light Microscopy platforms at i3S. Funds from Fundação para a Ciência e Tecnologia (FCT), co-financed by Fundo Europeu de Desenvolvimento Regional (FEDER) through Programa Operacional Competitividade e Internacionalização (POCI) (POCI-01-0145-FEDER-016390) to F. Janody and J. Paredes supported this work. The i3S Bioimaging and Advanced Light Microscopy scientific platforms are both members of the national infrastructure PPBI-Portuguese Platform of BioImaging (POCI-01-0145-FEDER-022122).

## Materials and Methods

### Generation of P-cad transgenic fly

The Gateway Cloning System (Life Technologies) was used to create pUASp-P-cad. The CDH3/P-cad coding sequence was first cloned into pENTR, and pUAS-P-cad was then generated by LR clonase II-mediated recombination of pENTR-PCAD into a modified Gateway vector pPW-attB [52]. The pUASp-PCAD transgene was then inserted into the *attP40* landing site by site-specific transgenesis (BestGene Inc).

### Fly strains and Genetics

Fly stocks used were *nub*-Gal4 [53], *ap*-Gal4; *eye*-Gal4, *GMR*-Gal4, *Sgs3-Gal4* (Bloomington #6870); *E-cad::GFP* [54], UAS-p35, *UAS-DE-cad-IR^HMS00693^, UAS-Src64B-IR^JF03234^,* UAS-*Src64B-IR^HMC03327^UAS-αPS2-IR^JF02695^, UAS-DMRTF-IR^JF02220^*, *UAS-DSRF-IR^JF02319^, tub-DMRTF.3XGFP* (Bloomington #58445); *UAS-DMRTF.ΔN; UAS-MRTF.S; bs^2^;* UAS-*MESK2-IR^JF03312^*; UAS-*det-IR^GL00572^*; UAS-*spire-IR^JF03233^*; UAS-*arpc2-IR^JF02845^*, UAS-*dia-IR^HM05027^*; UAS-*arp2-IR^JF02785^*, UAS-*arpc3A-IR^JF02370^*, UAS-*arpc3B-IR^JF02679^*, UAS-*arpc4-IR^JF01683^*, UAS-*arpc5-IR^JF03147^* (Bloomington Stock Center). All crosses were maintained at 25°C. Adult wings are from females only.

### Fly crosses

Fig. 1A

hsFLP; tub-FRT-stop-FRT-Gal4 UAS::GFP/CyO X UAS-P-cad

Fig. 1B

*nub*-Gal4, UAS-*P-cad* X DE-cad::GFP

Fig. 1C:

*nub*-Gal4; UAS-*mCD8-GFP* X w^1118^

*nub*-Gal4; UAS-*mCD8-GFP* X UAS-*DE-cad-IR^HMS00693^*

*nub*-Gal4; UAS-*P-cad* X UAS-*DE-cad-IR^HMS00693^*

Fig. 1D-F

*nub*-Gal4; UAS-*mCD8-GFP* X w^1118^

*nub*-Gal4, UAS-*P-cad* X UAS-*mCherry*

*nub*-Gal4, UAS-*P-cad* X UAS-*DE-cad-IR^HMS00693^*

Fig. 1G

*ap*-Gal4; UAS-*mCD8-GFP* X w^1118^

Fig. 1H

*ap*-Gal4; UAS-*mCD8-GFP* X UAS-*P-cad*

Fig. 1I

*nub*-Gal4; UAS-*mCD8-GFP* X w^1118^

*nub*-Gal4, UAS-*P-cad* X UAS-*mCherry*

*nub-Gal4; UAS-mCD8-GFP* X UAS-*Src64B-IR*^JF03234^ (*Src64B-IR #1*)

*nub*-Gal4, UAS-*P-cad* X UAS-*Src64B-IR*^JF03234^ (*Src64B-IR #1*)

*nub*-Gal4; UAS-*mCD8-GFP* X UAS-*Src64B*-IR^HMC03327^ (Src64B*-IR #2*)

*nub*-Gal4, UAS-*P-cad* X UAS-*Src64B*-IR^HMC03327^ (Src64B-IR #*2*)

*nub*-Gal4; UAS-*mCD8-GFP* X UAS-□*PS2-IR* ^JF02695^

*nub*-Gal4, UAS-*P-cad* X UAS-□*PS2-IR* ^JF02695^

Fig. 2A

*nub-Gal4; UAS-mCD8-GFP* X w^1118^

*nub*-Gal4, UAS-*P-cad* X UAS-*mCherry*

*nub*-Gal4; UAS-*mCD8-GFP* X UAS-*DMRTF-IR^JF02220^*

*nub*-Gal4, UAS-*P-cad* X UAS-*DMRTF-IR*^JF02220^

*nub*-Gal4; UAS-*mCD8-GFP* X UAS-*DSRF-IR*^JF02319^

*nub*-Gal4, UAS-*P-cad* X UAS-*DSRF-IR^JF02319^*

Fig. 2B

*nub*-Gal4; UAS-*mCD8-GFP* X w^1118^ (lane 1)

*nub*-Gal4, UAS-*P-cad* X UAS-*mCherry* (lane 2)

*nub*-Gal4, UAS-*P-cad* X UAS-*DMRTF-IR^JF02220^* (lane 3)

*nub*-Gal4, UAS-*P-cad* X UAS-*DSRF-IR^JF02319^* (lane 4)

Fig. 2C

*nub*-Gal4, UAS-*P-cad* X UAS-*mCherry*

*nub*-Gal4, UAS-*P-cad* X UAS-*DMRTF-IR*^JF02220^

*nub*-Gal4, UAS-*P-cad* X UAS-*DSRF-IR*^JF02319^

Fig. 2D

*tub*-Mrtf.3XGFP; *Sgs3*-Gal4 X UAS-*P-cad tub*-Mrtf.3XGFP X *Sgs3*-Gal4

Fig. 3A

*ap*-Gal4; UAS-*mCD8-GFP* X w^1118^

Fig. 3B

*ap*-Gal4; UAS-*mCD8-GFP* X UAS-*P-cad*

Fig. 3C

*nub*-Gal4; UAS-*mCD8-GFP* X w^1118^

*nub*-Gal4, UAS-*P-cad* X UAS-*mCherry*

*nub*-Gal4; UAS-*mCD8-GFP* X UAS-*arpc2-IR^JF02845^*

*nub*-Gal4, UAS-*P-cad* X UAS-*arpc2-IR^JF02845^*

*nub*-Gal4; UAS-*mCD8-GFP* X UAS-*spire-IR^JF03233^*

*nub*-Gal4, UAS-*P-cad* X UAS-*spire-IR^JF03233^*

*nub*-Gal4; UAS-*mCD8-GFP* X UAS-*dia-IR*^HM05027^

*nub*-Gal4, UAS-*P-cad* X UAS-*dia-IR*^HM05027^

Fig. 3D

*nub*-Gal4; UAS-*mCD8-GFP* X w^1118^ (lane 1)

*nub*-Gal4, UAS-*P-cad* X UAS-*mCherry* (lane 2)

*nub*-Gal4, UAS-*P-cad* X UAS-*arpc2-IR*^JF02845^ (lane 3)

*nub*-Gal4, UAS-*P-cad* X UAS-*spire-IR*^JF03233^ (lane 4)

Fig. 3E, F

*nub*-Gal4; UAS-*mCD8-GFP* X w^1118^ (lane 1)

*nub*-Gal4, UAS-*P-cad* X UAS-*mCherry* (lane 2)

*nub*-Gal4, UAS-*P-cad* X UAS-*dia-IR*^HM05027^ (lane 3)

Fig. 3F

*nub*-Gal4, UAS-*P-cad* X UAS-*mCherry*

*nub*-Gal4, UAS-*P-cad* X UAS-*dia-IR*^HM05027^

The FLPout system [55] was used to induce clonal P-cad overexpression (Fig. 1A). UAS-*P-cad* transgenic lines were crossed with y,w, hsFlp; tub-FRT-stop-FRT-Gal4 UAS:GFP/CyO and the progeny (y,w, hsFlp; UAS-P-cad/ tub-FRT-stop-FRT-Gal4, UAS:GFP) was heat-shocked to randomly induce Flippase-mediated removal of the FRT cassette.

### Cell culture conditions and drug treatments

The MCF10A-ER-Src cell line was kindly provided by K. Struhl [29]. Cells were grown in a humidified incubator at 37 °C, under a 5% CO_2_ atmosphere in Complete Growth Media (CGM), composed of DMEM/F12 growth media (Gibco, 11039-047), supplemented with 5% of Charcoal Stripped Horse Serum (Gibco, 16050-122), 20 ng/mL human EGF (Peprotech, AF-100-15), 0.5 μg/mL Hydrocortisone (Sigma, H0888), 100 ng/mL Cholera toxin from Vibrio cholerae (Sigma, C8052), 10 μg/mL Insulin (Sigma, I9278) and 0,5 μg/mL Puromycin (Merck, 540411). To treat cells with 4OH-TAM or EtOH, 50% confluent cells were plated and allowed to adhere for at least 24 hours before treatment with 1 μM 4OH-TAM (Merck, H7904) or with identical volume of EtOH for the indicated time. To assess the effect of preventing actin polymerization, cells were grown in CGM for 12 to 24 hours, washed and co-treated with 4OH-TAM or EtOH in the presence or absence of 0.5 μM LatA (Labclinics, 10010630) or of 40 μM CCG-203971 (Merck, SML1422) in restricted growth medium (RGM), composed of DMEM/F12, 0.5% CSHS, 0.5 μg/mL hydrocortisone, 100 ng/mL cholera toxin and 10 μg/mL insulin for the time points indicated.

### Real-time PCR analysis

RNAs were extracted using the NZY Total RNA Isolation Kit (NZYTech, MB13402). 0.5 μg of purified RNA samples were used for first strand cDNA synthesis (NZYTech, MB125) according to the manufacturer’s instructions. Real-time PCR was performed on 5 ng/μL cDNA using iTaq Universal SYBR Green Supermix (Bio-rad, 64361172) according to the manufacturer’s instructions. qRT-PCR was performed in triplicates using the CFX Touch Real Time detector system and relative fold change was calculated using ddCT method.

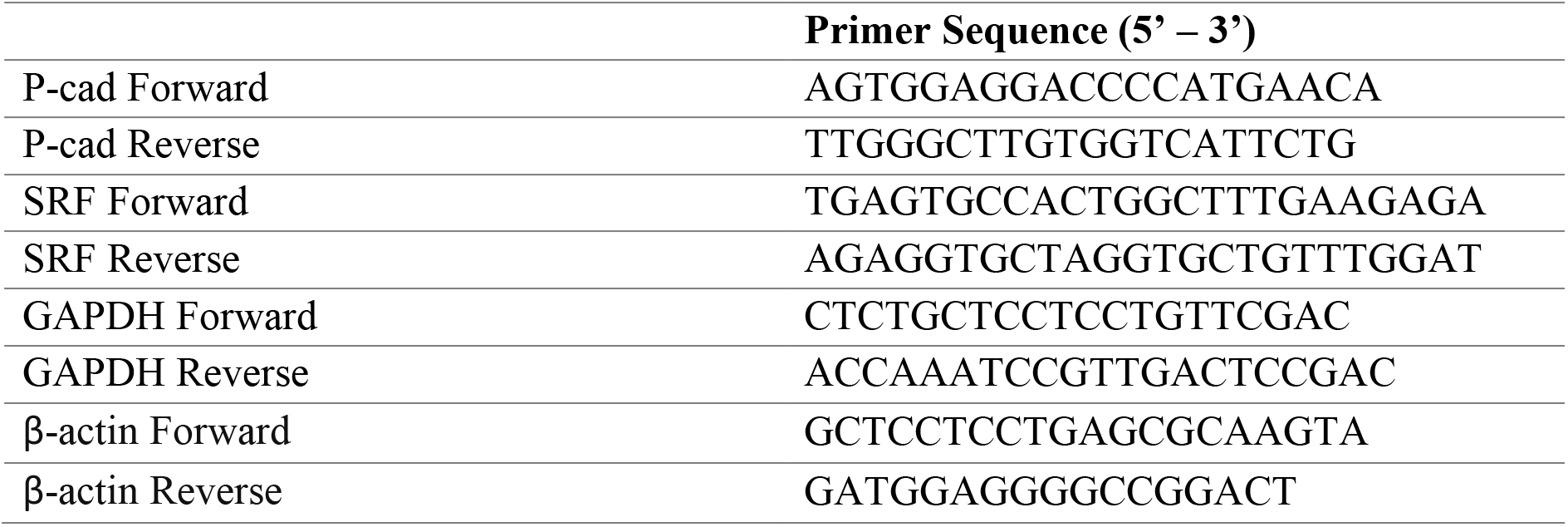

### Immunofluorescence Analysis

For immunofluorescence of the follicular epithelium, *Drosophila* ovaries were dissected in Schneider’s Insect Medium (Sigma-Aldrich, St. Louis, MI, USA) with 10% FBS and fixed in 4% paraformaldehyde. Following washing steps with 0.05% Tween-20 in PBS (PBS-T), and blocking with 10% BSA in PBS-T, ovaries were incubated overnight with Rabbit anti-Pcad (1:40; Cell Signaling; 2130S) and rat anti-E-cadherin (1:20, DSHB). Secondary antibodies used were Alexa Fluor 561 goat anti-mouse and the Alexa Fluor 647 goat anti-rabbit. For wing imaginal discs, P-cad and pSrc staining were performed by dissecting third instar larvae in phosphate buffer at pH 7 (0.1 M Na2HPO4, 0.1 M NaH2PO4 at a 72:28 ratio). Discs were then fixed in 4 % formaldehyde in PEM (0.1 M PIPES pH 7, 0.2 mM MgSO4, 1 mM EGTA) for 15 to 30 min, rinsed in phosphate buffer 0.2 % Triton for 15 min and incubated with rabbit anti-P-cad (1:50; Cell Signaling; 2130S) or anti-pSrc (pY419) (1:10; Invitrogen; 44-660G) overnight at 4°C. Discs were then rinsed 3 times 10 min in phosphate buffer 0.2% Triton, incubated for 1 hour at room temperature (RT) with secondary antibodies (Jackson ImmunoResearch) in phosphate buffer 0.2 % Triton supplemented with10 % horse serum and rinsed 3 more times 10 min before being mounted in Vectashield (Vector Labs, H-1000). For Phalloidin staining, discs were dissected and fixed as described above and incubated 1 hour at RT with Rhodamine-conjugated Phalloidin (Sigma, P-1951) at 0,3 mM in phosphate buffer 0.2 % Triton supplemented with 10 % horse serum. Discs were then rinsed 3 more times 10 min before being mounted in Vectashield (Vector Labs, H-1000). For salivary glands, third instar larvae were dissected on ice in phosphate buffer and fixed in 4 % formaldehyde in PEM, according to standard protocol. Samples were incubated with rabbit anti-P-cad (1:50; Cell Signaling, 2130S), mouse anti-GFP (1:500, DSHB-GFP-12A6) in phosphate buffer 0.2% Triton, supplemented with 10% FBS, overnight at 4°C. Alexa Fluor anti-rabbit and anti-mouse secondary antibodies were diluted in phosphate buffer 0.2% Triton, supplemented with 10% FBS (1:1000; AlexaFluor, Invitrogen). DAPI was used to counterstain the nuclei, and samples were mounted in 50% glycerol/PBS. Fluorescence images were obtained on LSM 510 Zeiss or Leica SP5 Live confocal microscopes using 10X dry and 40X objectives.

For MRTF-A nuclear translocation 75 000 MCF10A-ER-Src cells were seeded on poly-L-lysine-coated coverslips for at least 16 hours before being treated with 1 μM 4OH-TAM or identical volume of EtOH for 6 hours. Cells were then rinsed with PBS 1X and fixed with 4% paraformaldehyde in PBS 1X at pH 7 for 10 min at RT. Cells were then permeabilized with TBS - 0.1 % Triton (TBS-T) for 2 min at RT and blocked with blocking buffer (10 mM MES pH 6.1; 150 mM NaCl2; 5 mM EGTA pH 6.8; 5 mM MgCl2; 5 mM glucose; 2% FBS and 1 % BSA) for 1h at RT. Primary mouse anti-MRTF-A (G-8) monoclonal antibody (1:50; Santa Cruz, sc-390324) was incubated overnight at 4°C in blocking buffer. Coverslips were washed with PBS 1X three times for 5 min at RT and incubated with Donkey anti-mouse IgG Alexa Fluor 488 secondary antibody (1:200; Jackson ImmunoResearch, 715-095-150) and Rhodamine-conjugated Phalloidin (Sigma, P-1951) at 0.3 mM in blocking buffer for 1h at RT in dark. After three washes in PBS 1X, cells were stained with DAPI (1:500; Sigma D9542) for 5 min at RT, washed again with PBS 1X and mounted in Vectashield. Fluorescence images were obtained on a Leica SP5 confocal coupled to a Leica DMI6000, using the 63X 1.4 HCX PL APO CS Oil immersion objective. Image processing was performed using Imaris.

### Immunoblotting Analysis and Quantification

Protein extracts from wing imaginal discs were obtained by dissecting 6 discs in phosphate buffer at pH 7 (0.1M Na2HPO4, 0.1M NaH2PO4 at a 72:28 ratio). Discs in 28 μl PBS were lysed by adding 10 μl Laemmli Buffer 5X, 7 μl proteinase inhibitor 7X (Roche, 04693159001) and 5 μl phosphatase inhibitor 10X (Roche, 04906837001). Protein extracts from MCF10A-ER-Src were obtained by incubating cell pellets in Lysis Buffer SDS-Free containing 1% protease inhibitors and 1% phosphatase inhibitors on ice for 30 min. Lysis products were centrifuged at 4°C for 30 min at 14 000 rpm and protein was quantified using the Bradford method. Laemmli Buffer 5X was added to a final concentration of 1X. Protein extracts from wing discs and MCF10A-ER-Src cells were then boiled for 5 min at 95°C and centrifuged for 10 min at 8 000 rpm before being resolved by SDS-PAGE electrophoresis and transferred to a 0.45 μM PVDF blotting membrane (Amersham, 10600023), previously activated for 5 min in methanol. Membranes were blocked with 5% milk in TBS 0.1% Tween 20 and incubated with: mouse anti-P-cad (1:2500 BD, 610228), mouse anti-MRTF-A (1:1000 Santa Cruz, SC-390324), mouse anti-HSC70 (1:8000 Santa Cruz, SC-7298), or rabbit anti-H3 (1:300 Cell Signalling, 9715) diluted in TBS 0.1% Tween supplemented with 5 % non-fat milk or 3% BSA. Secondary antibodies used were HRP AP Donkey anti-mouse IgG (1:5000; Jackson Immunoresearch, 715-035-150) and HRP-conjugated AffiniPure Donkey anti-rabbit igG (1:5000; Jackson Immunoresearch, 711-035-152). Detection was performed using Immobilon Western Chemiluminescent HRP substrate (Milipore, P90719) and visualisation using ChemiDoc.

### Flow Cytometry

MCF10A-ER-Src cells treated with 4OH-TAM or EtOH for the indicated time were washed twice with Dulbecco’s phosphate buffered saline (DPBS, Biowest, L0615). Cells were then harvested with Versene Solution (Gibco, 15040-033), washed with PBS supplemented with 0.5% FBS (stain buffer), centrifuged at 1200 rpm for 5 min and resuspended in stain buffer. 500 000 cells were stained for 20 min at 4°C in the dark with the APC-conjugated anti-P-cad antibody (1:10; R&D system, FAB861A) and the Live/DeadTM Fixable Violet Dead Cell Stain Kit (1:1000; Invitrogen, L34955) in 100 μL of stain buffer. Cells were then washed twice with 1 mL of cold stain buffer followed by centrifugation at 1200 rpm for 5 min at 4 °C. Cells were resuspended in500 μL of cold stain buffer, passed through a 35 μm nylon cell strainer (Corning, 352235) to remove clumps and analysed on a FACS Canto II (BD Biosciences) equipped with the acquisition software BD FACSDiva (BD Biosciences). Data analysis was performed using FlowJo software version 10.5.3 (TreeStar, Inc.) and only viable (Live/Dead negative) single cells without debris were included into the analysis.

### Transfections and Dual Luciferase Renilla reporter assay

150 000 cells resuspended in 2 mL of CGM were plated in 6-well plates. After 12-24 hours, cells were transfected with 50 pmol of siCTR (Qiagen, 1027310), or siCDH3 (Qiagen, 1027416, GeneGlobe ID: SI02663941) and 0.720 μg of p3D.A-Luc and 1.44 μg of pRL-TK plasmids ^33^ using Lipofectamine 2000 (Invitrogen, 11668-019), according to the manufacturer’s instructions. After 12 hours, transfected cells were incubated for another 12 hours in RGM. Cells were then treated with EtOH or TAM in the presence or absence of LatA or CCG-203971 for 6 hours in RGM. After trypsinization with TrypLE^™^ Express (Gibco, 12604-021), cell pellets were resuspended in 250 μL of passive lysis buffer (PLB) 1X (Promega, E1941) and kept overnight at −20°C. Luciferase assay was performed using the Dual-Luciferase Reporter Assay system (Promega, PROME19600010) according to the manufacturer’s instructions. Briefly, 80 μL of lysed samples were added to 100 μL of luciferase assay reagent II (LARII) in a 96-well plate. Firefly luminescence was detected using synergy Mx. Then, 100 μL of 1X Stop and Glo solution were added to detect Renilla luminescence. Each sample was evaluated in triplicates.

### Trypan-blue cell viability assay

Cells were washed with PBS and trypsinized using TrypLE™ Express for 15 min in a 5% CO2 humidified incubator at 37°C. After centrifugation at 200 g for 5 min, cell pellets were resuspended in 20 μL RGM. For each experimental condition, 5 μL of cell suspension in the same volume of Trypan-Blue (Lonza, LONZ17-942E) were counted in a Neubauer chamber.

### Cell Cycle Profile

265 000 cells in 4 mL of CGM were plated in a T25 flask. After 12-24 hours, cells were washed three times with DMEM/F12 and incubated for 12 hours in RGM. Cells were then treated with EtOH or TAM in the presence or absence of 40 μM CCG-203971 for 12, 24 or 36 hours in RGM, trypsinized with TrypLE^™^ Express and collected in 5 mL round bottom polystyrene tubes (Corning, 352235). After centrifugation for 5 min at 1000 rpm at 4°C, cells were resuspended in 1 mL of FACS Buffer 1, composed of 2% Fetal Bovine Serum heat-inactivated (Biowest, S181BH) in PBS1X, centrifuged again at 4°C for 5 min at 1000 rpm and resuspended in 500μL FACS Buffer 1. Cells were then fixed with 1.5 mL 70% Ethanol, added drop-by-drop, while gently vortexed, followed by a 30 min incubation at 4°C. After 5 min centrifugation at 2000 rpm at 4°C, cell were resuspended in 3 mL PBS1X, incubated 30 min on ice, pelleted again by centrifugation for 5 min at 2000 rpm at 4°C and resuspended in 300μL of FACS Buffer 2, composed of 100 μg/mL RNAse A (Qiagen, 19101) and 20 μg/mL Propidium Iodide (Sigma, P4170) in PBS1X. Samples were incubated for 30 min in a 37°C water bath in the dark.

Flow Cytometry was performed at low flow rate using a BD Accuri C6 Flow Cytometer (Becton-Dickinson, Franklin Lakes, NJ, USA). Cell Cycle profile was analysed using FlowJo 10.7.1 software (Tree Star, Inc., Ashland, OR, USA), Cell Cycle platform, using the Watson Model.

### Invasion assay in collagen

20 000 cells in 200 μL of CGM were plated in a 96-well plate, previously coated with 30μL 0.7% agarose (Lonza, 50004). Cells were allowed to form spheroids for 24 hours. On the next day, medium was replaced by 60 μL of 5 mg/mL Collagen Type I (VWR, 734-1085) in 1M NaOH and 1X DMEM/F12. After 1 hour, spheroids were treated with 4 μM TAM or the same volume of EtOH in the presence or absence of 160 μM CCG-203971 and incubated in a 5% CO2 humidified incubator at 37°C for 36 hours. Brightfield pictures were taken using the 5X objective on a brightfield microscope.

### Mammosphere assay

150 000 cells in 2 mL of CGM were plated in a 6-well plate. After 12-24 hours, cells were treated with EtOH or TAM in the presence or absence of 40 μM CCG-203971 for 36 hours in RGM. Cells were trypsinized with Versene, resuspended in 1 mL of PBS 1X and passed three times through a 25 gauge needle to obtain single cell suspensions. 10 000 cells were then plated in 6-well plates previously coated with Poly(2-hydrocyethyl methacrylate) (PolyHEMA) and incubated in 2 mL of mammosphere medium composed of 1:1 DMEM/F12, 20 ng/mL human EGF, 40 μg/mL Insulin, 500 ng/mL Hydrocortisone, 1 % Penincilin/Streptomycin (Merck, 15070-063) and 1:1 Supplement B-27 Minus Vitamine A (Gibco,12587-010), filtered with a 0.45 μm filter (VWR, 514-4127). 6 days after incubation in a 5% CO2 humidified incubator at 37°C, brightfield pictures were acquired using the 5X objective on a brightfield microscope. Each condition was evaluated in triplicates.

### Quantifications

The NIH Image J program was used to quantify wing disc area. Except for *nub>GFP* control wings, comparisons were performed on wings containing the same number of UAS transgenes. UAS-*GFP* or UAS-*mCherry* were used for normalisation when needed. Each disc was outlined and measured using the *Area* function, which evaluates size in square pixels. To quantify the ratio of the *nub>GFP* domain over the total wing disc area, the ratio between the area of the GFP domain and the area of the whole disc domain, measured using the *Area* function for each disc, was performed. To evaluate nuclear accumulation of MRTF.3XGFP, we used surface segmentation tools from Imaris Software to define nuclei area, on 3D projections of salivary gland images. The intensity of MRTF.3XGFP signals within the defined nuclei was extracted automatically. The NIH Image J program was used to evaluate MRTF-A nuclear accumulation. Nuclear MRTF-A fluorescence intensity was measured as mean grey intensity value for each nucleus, defined using the DAPI channel. Fold changes in nuclear MRTF-A was calculated after normalisation to MRTF-A nuclear intensity in EtOH-treated cells.

Relative luciferase activity was quantified by normalising Firefly Luciferase activity to their respective Renilla Luciferase activity. Western-blot quantification was performed using the Image Lab software. Briefly, each band was selected using the tab “lanes and bands” function and adjusted for noise using the “lane profile” function. Band intensities were determined in the “Analysis table” function. Spheroid’s circularity was quantified using the NIH Image J program. Each spheroid was outlined using the freehand selection tool and measured using the shape descriptor tool. Mammospheres with ≥ 60 μm diameter were counted using the 5X objective on a brightfield microscope. To calculate mammosphere forming efficiency, the number of mammospheres counted was divided by the number of cells plated (10 000 cells). GraphPad Prism (9.0) was used for data presentation and statistical analysis.

