## Supplementary Figures for "Activation of the actin/MRTF-A/SRF signalling pathway in pre-malignant mammary epithelial cells by P-cadherin is essential for transformation"

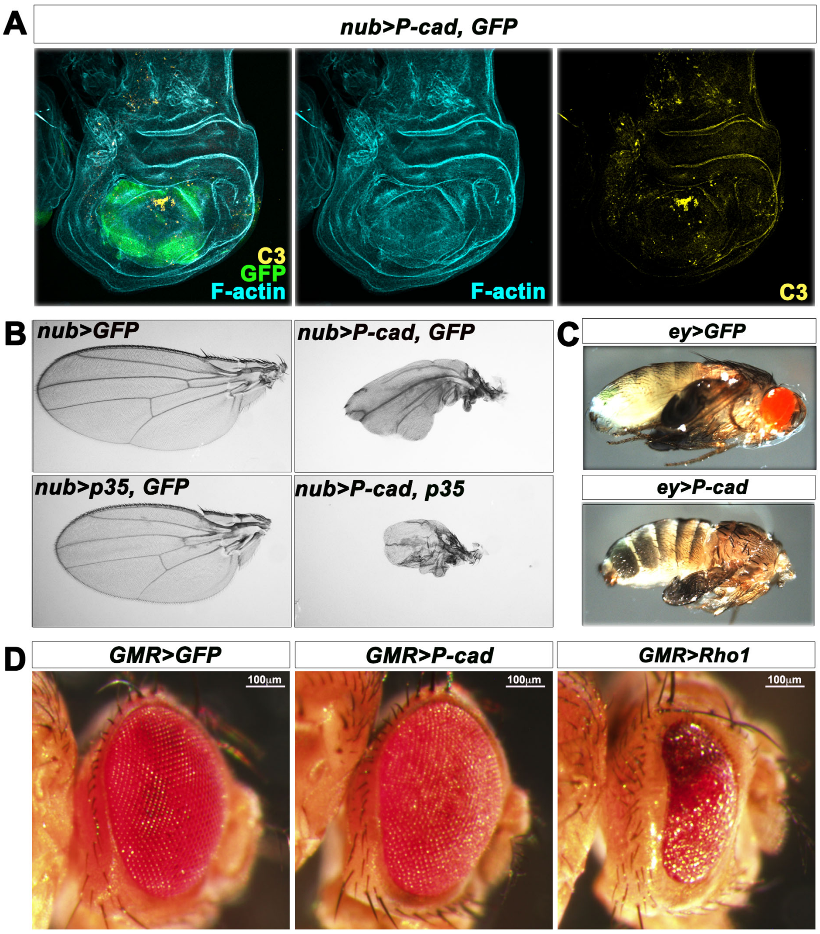


**Supplementary Figure 1: P-cad expression affects dividing epithelia without triggering massive apoptosis. (A)** Standard confocal sections of third instar wing imaginal discs expressing UAS-*mCD8-GFP* (green) and UAS-*P-cad* under *nub*-Gal4 control and stained with Phalloidin (cyan blue) to mark F-actin and anti-activated Caspase 3 (C3) (yellow). **(B)** Adult wings in which *nub*-Gal4 drives UAS-*mCD8-GFP* alone or UAS-*mCD8-GFP* and UAS-*P-cad* or UAS-*mCD8-GFP* and UAS-*p35* or UAS-*p35* and UAS-*P-cad.* **(C)** Pupae in which *ey*-Gal4 drives UAS-*mCD8-GFP* alone or UAS-*P-cad.* **(D)** Adult eyes in which *GMR*-Gal4 drives UAS-*mCD8-GFP* alone or UAS-*P-cad* or UAS-*Rho1*.

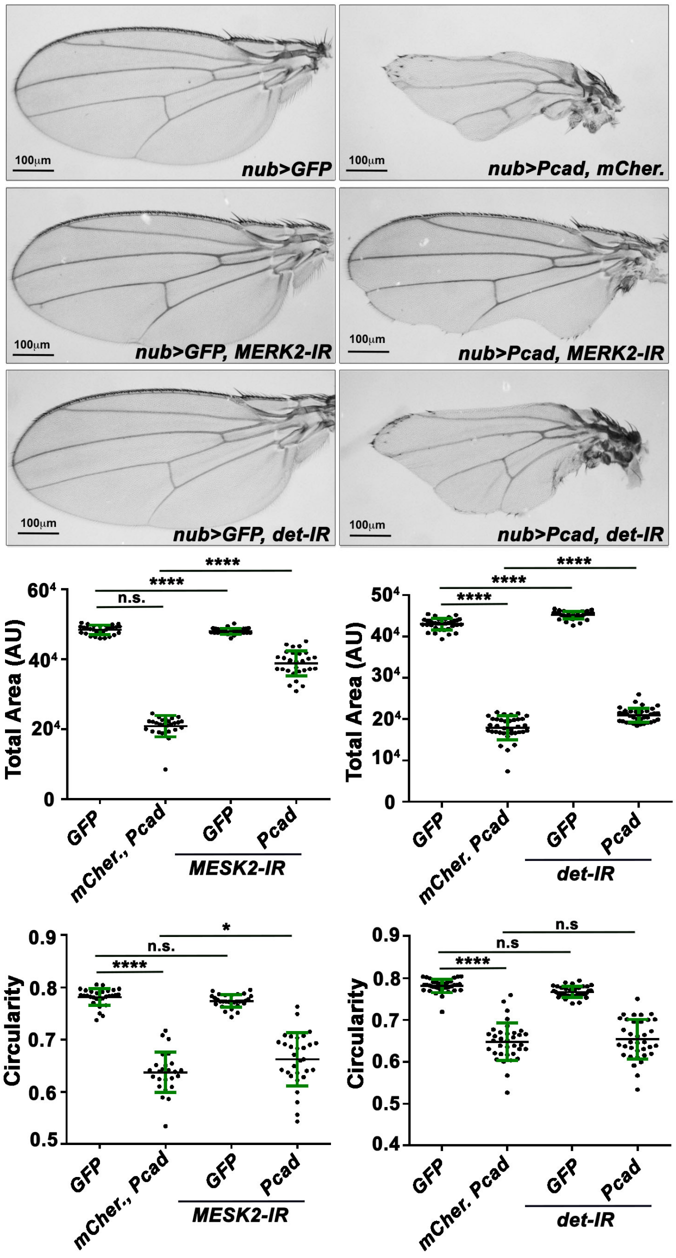


**Supplementary Figure 2: Knocking down the DMRTF-DSRF transcriptional targets *MESK2* or *det* suppresses the *P-cad*-expressing wing phenotype.** (Top panels) Adult wings in which *nub*-Gal4 drives UAS-*mCD8-GFP* or UAS-*P-cad* and UAS-*mCherry* or UAS-*MESK2-IR*$^{JF03312}$ and UAS-*mCD8-GFP* or UAS-*MESK2-IR*$^{JF03312}$ and UAS-*P-cad* or UAS-*det-IR*$^{GL00572}$ and UAS-*mCD8-GFP* or UAS-*det-IR*$^{GL00572}$ and UAS-*P-cad.* (Bottom panels) Quantifications of the total area or circularity of wings for the genotypes indicated. Error bars indicate SD; n.s. indicate non-significant; * indicates P<0.05; **** indicates P<0.0001**.** Statistical significance was calculated using one-way ANOVA tests.

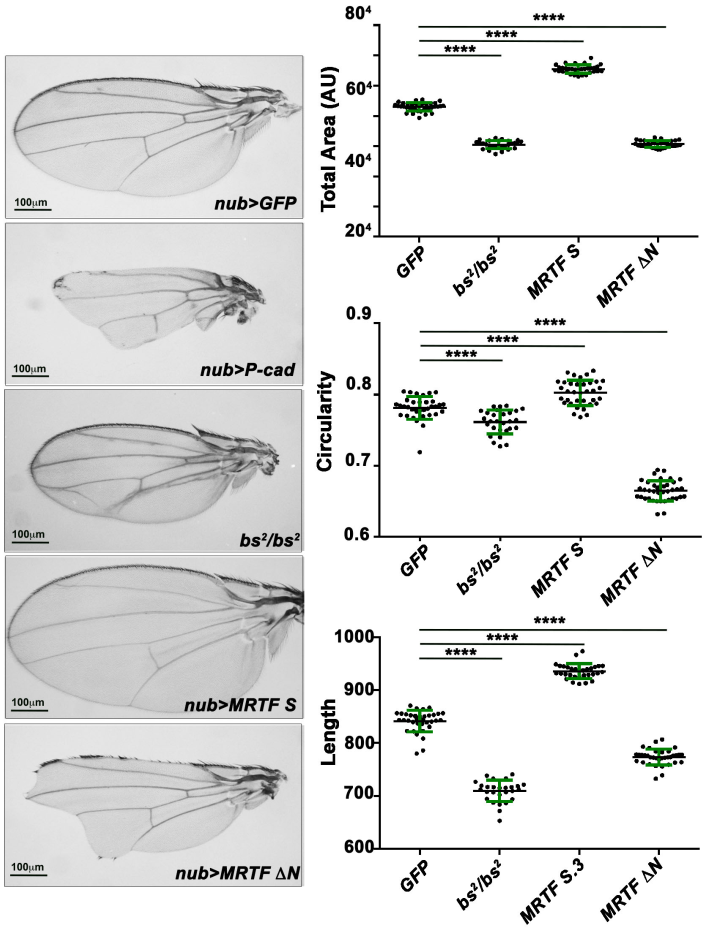


**Supplementary Figure 3: Expressing a constitutive active form of *DMRTF* phenocopies the *P-cad*-expressing wing phenotype.** (Left panels) Adult wings in which *nub*-Gal4 drives UAS-*mCD8-GFP* or UAS-*P-cad* and UAS-*mCherry* or UAS-*DMRTF* or UAS-$DMRTF^{\Delta N}$ or homozygous mutant for the $DSRF^2$ allele ($DSRF^2/DSRF^2$). (Right panels) Quantifications of the total area or circularity or anterior-posterior (A/P) length of adult wings for the genotypes indicated. Error bars indicate SD; **** indicates P<0.0001**.** Statistical significance was calculated using one-way ANOVA tests.

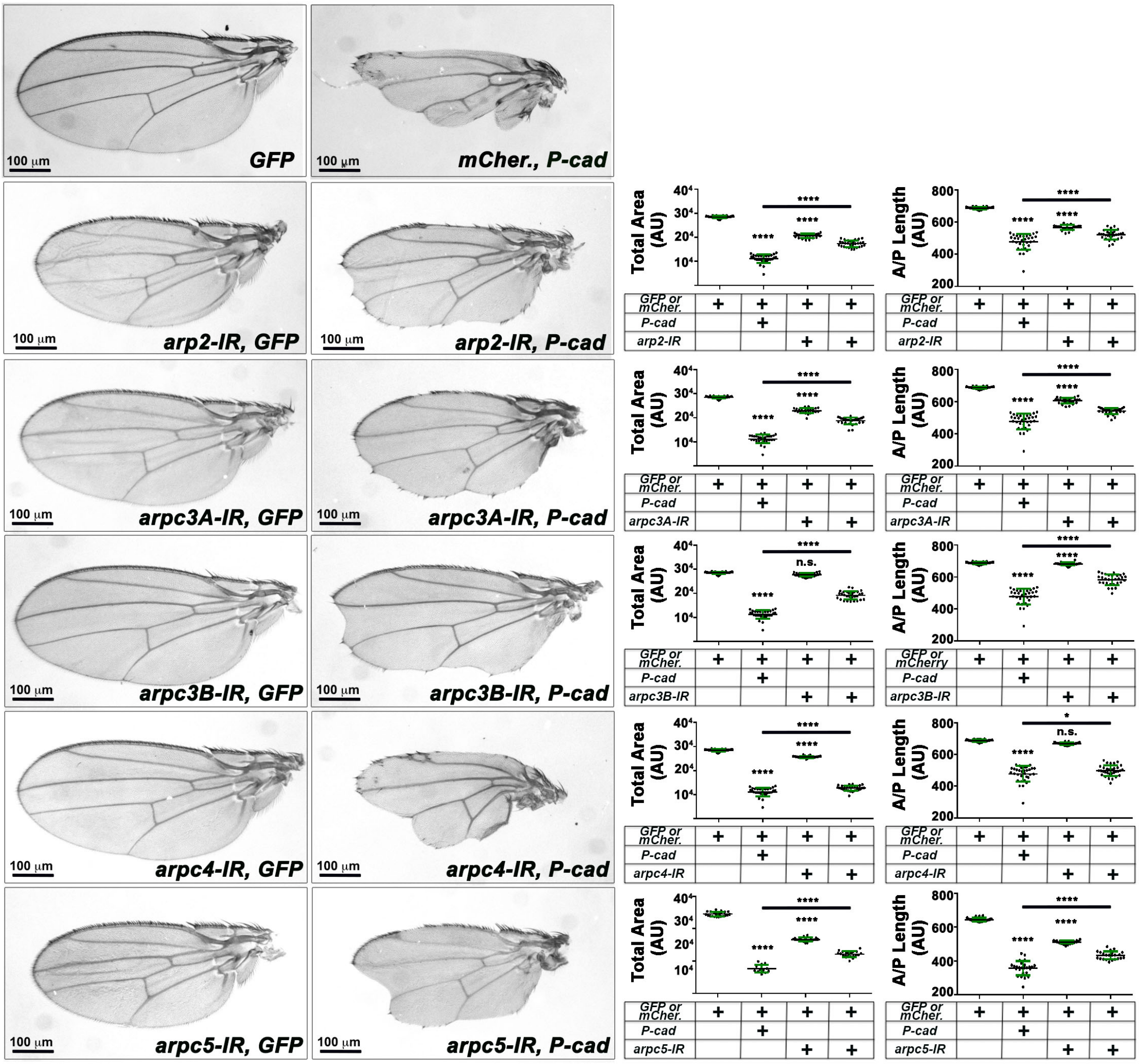

**Supplementary Figure 4: Knocking down subunits of the Arp2/3 complex suppresses the *P-cad*-expressing wing phenotype.** (Left panels) Adult wings in which *nub*-Gal4 drives UAS-*mCD8-GFP* or UAS-*P-cad* and UAS-*mCherry* or UAS-*arp2-IR*$^{JF02785}$ and UAS-*mCD8-GFP* or UAS-*arp2-IR*$^{JF02785}$ and UAS-*P-cad* or UAS-*arpc3A-IR*$^{JF02370}$ and UAS-*mCD8-GFP* or UAS-*arpc3A-IR*$^{JF02370}$ and UAS-*P-cad* or UAS-*arpc3B-IR*$^{JF02679}$ and UAS-*mCD8-GFP* or UAS-*arpc3B-IR*$^{JF02679}$ and UAS-*P-cad* or UAS-*arpc4-IR*$^{JF01683}$ and UAS-*mCD8-GFP* or UAS-*arpc4-IR*$^{JF01683}$ and UAS-*P-cad* or UAS-*arpc5-IR*$^{JF03147}$ and UAS-*mCD8-GFP* or UAS-*arpc5-IR*$^{JF03147}$ and UAS-*P-cad.* (Right panels) Quantifications of the total area or anterior-posterior (A/P) length of adult wings for the genotypes indicated. Error bars indicate SD; n.s. indicate non-significant; * indicates P<0.05; **** indicates P<0.0001**.** Statistical significance was calculated using one-way ANOVA tests.

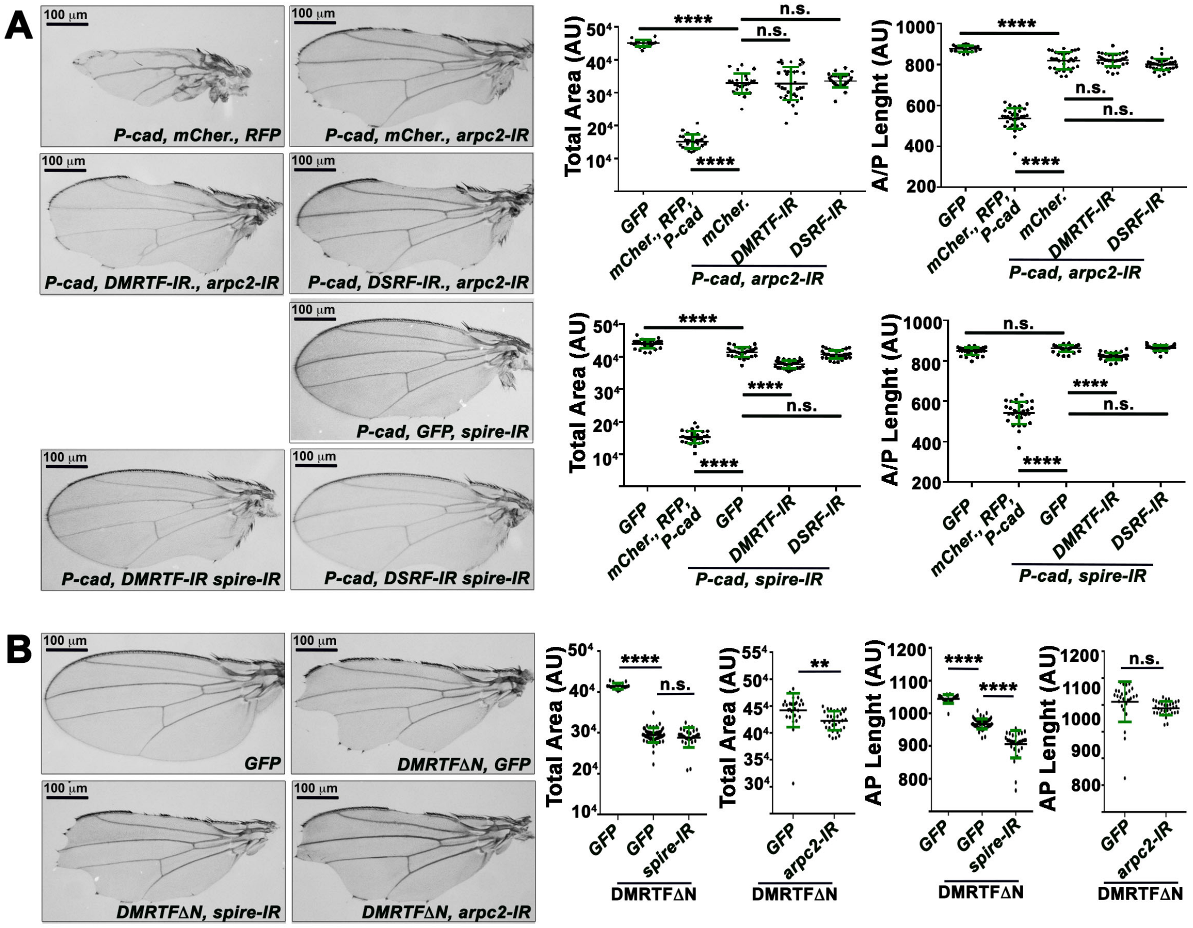


**Supplementary Figure 5: *arpc2* and *spire* are required downstream of *P-cad* and upstream or in parallel of $DMTF^{\Delta N}$ to affect wing differentiation. (A)** (Left panels) Adult wings in which *nub*-Gal4 drives UAS-*P-cad*, UAS-*mCher.* and UAS-*RFP* or UAS-*P-cad*, UAS-*mCher.* and UAS-*arpc2-IR*$^{JF02845}$ or UAS-*P-cad*, UAS-*arpc2-IR*$^{JF02845}$ and UAS-*DMRTF-IR*$^{JF02220}$ or UAS-*P-cad*, UAS-*arpc2-IR*$^{JF02845}$ and UAS-*DSRF-IR*$^{JF02319}$ or UAS-*P-cad*, UAS-*mCher.* and UAS-*spire-IR*$^{JF03233}$ or UAS-*P-cad*, UAS-*spire-IR*$^{JF03233}$ and UAS-*DMRTF-IR*$^{JF02220}$ or UAS-*P-cad*, UAS-*spire-IR*$^{JF03233}$ and UAS-*DSRF-IR*$^{JF02319}$. (Right panels) Quantifications of the total area or anterior-posterior (A/P) length of adult wings for the genotypes indicated. **(B)** (Left panels) Adult wings in which *nub*-Gal4 drives UAS-*mCD8-GFP* or UAS-*mCD8-GFP* and UAS-$DMRTF^{\Delta N}$ or UAS-$DMRTF^{\Delta N}$ and UAS-*spire-IR*$^{JF03233}$ or UAS-$DMRTF^{\Delta N}$ and UAS-*arpc2-IR*$^{JF02845}$. (Right panels) Quantifications of the total area or anterior-posterior (A/P) length of adult wings for the genotypes indicated. Error bars indicate SD. n.s. indicate non-significant; ** indicates P<0.01; **** indicates P<0.0001**.** Statistical significance was calculated using one-way ANOVA tests.
